# Hypoxia-inducible factor induces cysteine dioxygenase and promotes cysteine homeostasis in *Caenorhabditis elegans*

**DOI:** 10.1101/2023.05.04.538701

**Authors:** Kurt Warnhoff, Sushila Bhattacharya, Jennifer Snoozy, Peter C. Breen, Gary Ruvkun

**Affiliations:** Pediatrics and Rare Diseases Group, Sanford Research, Sioux Falls, SD 57104, USA; Department of Pediatrics, Sanford School of Medicine, University of South Dakota, Sioux Falls, SD 57105 USA; Department of Molecular Biology, Massachusetts General Hospital, Boston, MA 02114, USA

## Abstract

Dedicated genetic pathways regulate cysteine homeostasis. For example, high levels of cysteine activate cysteine dioxygenase, a key enzyme in cysteine catabolism in most animal and many fungal species. The mechanism by which cysteine dioxygenase is regulated is largely unknown. In an unbiased genetic screen for mutations that activate cysteine dioxygenase (*cdo-1*) in the nematode *C. elegans,* we isolated loss-of-function mutations in *rhy-1* and *egl-9,* which encode proteins that negatively regulate the stability or activity of the oxygen-sensing hypoxia inducible transcription factor (*hif-1*). EGL-9 and HIF-1 are core members of the conserved eukaryotic hypoxia response. However, we demonstrate that the mechanism of HIF-1-mediated induction of *cdo-1* is largely independent of EGL-9 prolyl hydroxylase activity and the von Hippel-Lindau E3 ubiquitin ligase, the classical hypoxia signaling pathway components. We demonstrate that *C. elegans cdo-1* is transcriptionally activated by high levels of cysteine and *hif-1*. *hif-1-*dependent activation of *cdo-1* occurs downstream of an H_2_S-sensing pathway that includes *rhy-1, cysl-1,* and *egl-9. cdo-1* transcription is primarily activated in the hypodermis where it is also sufficient to drive sulfur amino acid metabolism. Thus, the regulation of *cdo-1* by *hif-1* reveals a negative feedback loop that maintains cysteine homeostasis. High levels of cysteine stimulate the production of an H_2_S signal. H_2_S then acts through the *rhy-1/cysl-1/egl-9* signaling pathway to increase HIF-1-mediated transcription of *cdo-1,* promoting degradation of cysteine via CDO-1.

## Introduction

Cysteine is a sulfur-containing amino acid that mediates many oxidation/reduction reactions of proteins, is the redox center of the abundant antioxidant tripeptide glutathione which also serves as a major cysteine reserve, and is essential for iron-sulfur cluster assembly in the mitochondrion (1, 2). Cysteine residues in many proteins are in close proximity in the primary or folded protein sequence and are oxidized in the endoplasmic reticulum to form intra- and interprotein disulfide linkages, most commonly in secreted proteins which mediate intercellular signaling and defense (3–5). In many enzymes, the reactivity of the cysteine sulfur is key for the coordination of metals such as zinc or iron, which support protein structure and catalytic activity (6–8). Cysteine is also a key source of hydrogen sulfide (H_2_S), a volatile signaling molecule (9). While cysteine has these essential functions, excess cysteine is also toxic. High levels of cysteine impair mitochondrial respiration by disrupting iron homeostasis (10), acts as a neural excitotoxin (11), and promotes the formation of toxic levels of hydrogen sulfide gas (9, 12, 13). Given this balance between essential and toxic, cysteine homeostasis is key for the health of cells and organisms.

CDO1-mediated oxidation is the primary pathway of cysteine catabolism when sulfur amino acid (methionine or cysteine) availability is normal or high (14). The dipeptide cystathionine is a key intermediate in this pathway. Cystathionine is catabolized by cystathionase (CTH-2 in *C. elegans,* CTH in mammals) producing cysteine and α-ketobutyrate. Cysteine is further oxidized using dissolved atmospheric dioxygen to cysteinesulfinate by cysteine dioxygenase (CDO-1 in *C. elegans,* CDO1 in mammals) (**Fig. 1A**) (15). The further oxidation of cysteinesulfinate downstream of CDO-1 generates highly toxic sulfites that are normally oxidized to more benign sulfate by sulfite oxidase (**Fig. 1A**).

**Figure 1:**
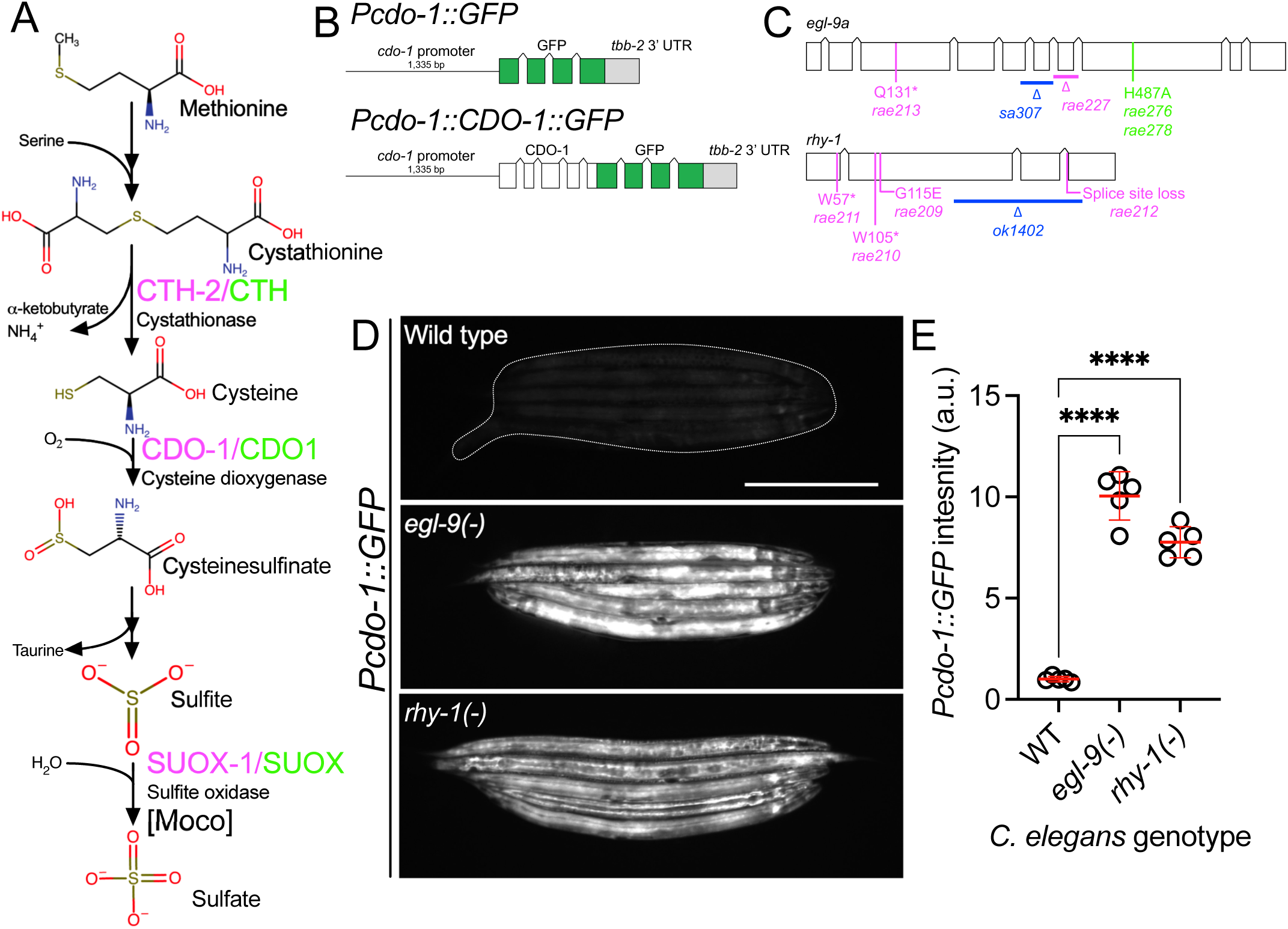
*egl-9* and *rhy-1* inhibit *cdo-1* transcription. A) Pathway for sulfur amino acid metabolism beginning with methionine. We highlight the roles of cystathionase (CTH-2/CTH), cysteine dioxygenase (CDO-1/CDO1), and the Moco-requiring sulfite oxidase enzyme (SUOX-1/SUOX). *C. elegans* enzymes (magenta) and their human homologs (green) are displayed. B) *Pcdo-1::GFP* promoter fusion (upper) and *Pcdo-1::CDO-1::GFP* C-terminal protein fusion (lower) transgenes used in this work are displayed. Boxes indicate exons, connecting lines indicate introns. The *cdo-1* promoter is shown as a straight line. C) *egl-9a* and *rhy-1* gene structures. Boxes indicate exons and connecting lines are introns. Colored annotations indicate mutations generated or used in our work. Magenta; chemically-induced mutations that activated *Pcdo-1::CDO-1::GFP* fusion protein. Blue; reference null alleles isolated independent of our work. Green; CRISPR/Cas9-generated mutation that inactivates the prolyl hydroxylase domain of EGL-9. D) Expression of *Pcdo-1::GFP* transgene is displayed for wild-type, *egl-9(sa307),* and *rhy-1(ok1402) C. elegans* animals at the L4 stage. Scale bar is 250μm. White dotted line outlines animals with basal GFP expression. For GFP imaging, exposure time was 100ms. E) Quantification of GFP expression displayed in Fig. 1D. Individual datapoints are shown (circles) as are the mean and standard deviation (red lines). *n* is 5 individuals per genotype. Data are normalized so that wild-type expression of *Pcdo-1::GFP* is 1 arbitrary unit (a.u.). ****, p<0.0001, ordinary one-way ANOVA with Dunnett’s post hoc analysis.

As a critical player in cysteine homeostasis, CDO1 is a highly regulated enzyme: the activity and abundance of CDO1 increase dramatically in cells and animals fed excess cysteine and methionine (16). CDO1 levels are governed both transcriptionally and post-translationally via proteasomal degradation (17–21). However, the molecular players that sense high levels of cysteine and promote CDO1 activation remain incompletely defined.

Understanding these regulatory mechanisms is an important goal as CDO1 dysfunction is implicated in disease. CDO1 is a tumor suppressor whose activity is silenced in diverse cancers via promoter methylation, a potential biomarker of tumor grade and progression (22–24). Decreased CDO1 activity may support tumor cell growth by reducing reactive oxygen species and decreasing drug susceptibility (25–28).

Using the nematode *C. elegans,* we have previously shown that CDO-1 is a key player in the pathophysiology of two fatal inborn errors of metabolism; isolated sulfite oxidase deficiency (ISOD) and molybdenum cofactor deficiency (MoCD) (29, 30). Molybdenum cofactor (Moco) is an essential 520 Dalton prosthetic group synthesized from GTP by a conserved multistep biosynthetic pathway that is present in about 2/3 of bacterial genomes and nearly all eukaryotic genomes (31, 32). *C. elegans* can either retrieve Moco synthesized by the bacteria it consumes or can synthesize Moco *de novo* using its own Moco biosynthetic pathway (33). *C. elegans* strains carrying mutations in genes encoding Moco biosynthetic enzymes (*moc*) and feeding on wild-type *E. coli* develop normally. Yet, these same *moc-*mutant *C. elegans* when fed on Moco-deficient *E. coli* as their sole nutritional source are inviable, arresting development at an early larval stage. A saturated genetic selection for mutations that suppress this inviability identified multiple independent mutations in *cth-2* or *cdo-1*, genes which encode enzymes in the cysteine biosynthetic and degradation pathway. The toxic sulfites produced downstream of CDO-1 are normally oxidized to more benign sulfate by Moco-requiring sulfite oxidase (SUOX-1 in *C. elegans,* SUOX in mammals) an essential enzyme in *C. elegans* and humans (**Fig. 1A**) (30, 33). Therefore, loss of *cdo-1* or *cth-2* suppresses the lethality caused by both Moco and sulfite oxidase deficiencies in *C. elegans* by preventing the production of toxic sulfites (33–35). Thus, understanding the fundamental mechanisms that govern the levels and activity of CDO-1 is critical to generating new therapeutic hypotheses to treat these diseases.

To define genes that regulate *cdo-1* levels and activity, we performed an unbiased genetic screen in the nematode *C. elegans* to identify mutations that increase the expression or abundance of a *Pcdo-1::CDO-1::GFP* reporter transgene. We identified multiple independent loss-of-function mutations in two genes, *egl-9* and *rhy-1*, that dramatically increase expression of this P*cdo-1::CDO-1::GFP* transgene. These enzymes act in the hypoxia and H_2_S-sensing pathway, and we demonstrate that the conserved hypoxia-inducible transcription factor (HIF-1) activates *cdo-1* transcription in this pathway (36–38). We further show that high levels of cysteine promote *cdo-1* transcription, and that *hif-1* and *cysl-1* (another component of the H_2_S-sensing pathway) are required for viability under high cysteine conditions. We demonstrate that transcriptional activation of *cdo-1* via HIF-1 promotes CDO-1 activity and establish the *C. elegans* hypodermis as a key tissue of CDO-1 activation and function. Unexpectedly, we find that *cdo-1* regulation is governed by a HIF-1 pathway largely independent of EGL-9 prolyl hydroxylase activity and von Hippel-Lindau (VHL-1), the canonical O_2_ -sensing pathway (39, 40). These data establish a new connection between the HIF-1/H_2_S-sensing pathway and sulfur amino acid catabolism governed by CDO-1.

## Results

### *egl-9* and *rhy-1* negatively regulate *cdo-1* transcription

To identify regulators of CDO-1 expression or activity, we engineered a transgene expressing a C-terminal green fluorescent protein (GFP) fusion to the full-length CDO-1 protein driven by the native *cdo-1* promoter (*Pcdo-1::CDO-1::GFP*, **Fig. 1B**). Transgenic animals were generated by integrating the *Pcdo-1::CDO-1::GFP* fusion protein into the *C. elegans* genome (41). The *Pcdo-1::CDO-1::GFP* fusion protein was functional and rescued a *cdo-1* loss of function mutation: the *Pcdo-1::CDO-1::GFP* fusion protein reverses the suppression of Moco-deficient lethality caused by *cdo-1* loss of function (**Fig. S1**). The reanimation of Moco-deficient lethality by the transgene depends on CDO-1 enzymatic activity because a transgene expressing an active-site mutant *Pcdo-1::CDO-1[C85Y]::GFP* does not rescue the *cdo-1* mutant phenotype (**Fig. S1**) (33, 42). Thus, the *Pcdo-1::CDO-1::GFP* transgenic fusion protein is functional, suggesting that its expression pattern reflects endogenous protein expression, localization, and levels.

We performed a mutagenesis screen to identify genes that control the expression or accumulation of CDO-1 protein. Specifically, we performed an EMS chemical mutagenesis of *C. elegans* and screened in the F2 generation, after newly induced random mutations were allowed to become homozygous, for mutations that caused increased GFP accumulation by the *Pcdo-1::CDO-1::GFP* reporter transgene (43). Using whole-genome sequencing, we determined that two independently isolated mutants carried distinct mutations in *egl-9* and four other independently isolated mutants had unique mutations in *rhy-1* (**Fig. 1C)**. The presence of multiple independent alleles suggests these mutations in *egl-9* or *rhy-1* are causative for the increased *Pcdo-1::CDO-1::GFP* expression or accumulation observed. *egl-9* encodes the O_2_ -sensing prolyl hydroxylase and orthologue of mammalian EglN1. *rhy-1* encodes the regulator of <u>hy</u>poxia inducible transcription factor, an enzyme with homology to membrane-bound O-acyltransferases (36, 39). The genetic screen produced nonsense alleles of both *egl-9* and *rhy-1* suggesting the increased *Pcdo-1::CDO-1::GFP* expression or accumulation is caused by loss of *egl-9* or *rhy-1* function (**Fig. 1C**). Inactivation of *egl-9* or *rhy-1* activates a transcriptional program mediated by the hypoxia inducible transcription factor, HIF-1 (36, 39). To determine if *egl-9* or *rhy-1* regulate *cdo-1* transcription, we engineered a separate reporter construct where only GFP (rather than the full length CDO-1::GFP fusion protein) is transcribed by the *cdo-1* promoter (*Pcdo-1::GFP,* **Fig. 1B**). This *Pcdo-1::GFP* transgene was introduced into strains with independently isolated *egl-9(sa307)* and *rhy-1(ok1402)* null reference alleles (36, 44). The *egl-9(sa307)* and *rhy-1(ok1402)* mutations dramatically induce expression of GFP driven by the *Pcdo-1::GFP* transcriptional reporter transgene (**Fig. 1D,E**). The activation of *cdo-1* transcription by independently isolated *egl-9* or *rhy-1* mutations demonstrates that the *egl-9* and *rhy-1* mutations isolated in our screen for the induction or accumulation of *Pcdo-1::CDO-1::GFP* are the causative genetic lesions. Furthermore, these data demonstrate that *egl-9* and *rhy-1* are necessary for the normal transcriptional repression of *cdo-1*.

### HIF-1 directly activates *cdo-1* transcription downstream of the *rhy-1, cysl-1, egl-9* genetic pathway

*rhy-1* and *egl-9* act in a pathway that regulates the abundance and activity of the HIF-1 transcription factor (36, 38). The activity of *rhy-1* is most upstream in the pathway and negatively regulates the activity of *cysl-1,* which encodes a cysteine synthase-like enzyme of probable algal origin (45). CYSL-1 directly binds to and inhibits EGL-9 in an H_2_S-modulated manner (38). EGL-9 uses molecular oxygen as well as an α-ketoglutarate cofactor to directly inhibit HIF-1 via prolyl hydroxylation, which recruits the VHL-1 ubiquitin ligase to ubiquitinate HIF-1, targeting it for degradation by the proteasome **(Fig. 5A)** (46–48). Given that loss-of-function mutations in *rhy-1* or *egl-9* activate *cdo-1* transcription, we tested if *cdo-1* transcription is activated by HIF-1 as an output of this hypoxia/H_2_S-sensing pathway. We performed epistasis studies using null mutations that inhibit the activity of *hif-1* (*cysl-1(ok762)* and *hif-1(ia4)*) or activate *hif-1* (*rhy-1(ok1402)* and *egl-9(sa307)*). The induction of *Pcdo-1::GFP* by *egl-9* inactivation was dependent upon the activity of *hif-1,* but not on the activity of *cysl-1* (**Fig. 2A,B)**. In contrast, induction of *Pcdo-1::GFP* by *rhy-1* inactivation was dependent upon the activity of both *hif-1* and *cysl-1* (**Fig. 2A,C)**. These results reveal a genetic pathway whereby *rhy-1, cysl-1,* and *egl-9* function in a negative-regulatory cascade to control the activity of HIF-1 which transcriptionally activates *cdo-1.* Our epistasis studies of *cdo-1* transcriptional regulation by HIF-1 align well with previous analyses of this genetic pathway in the context of transcription of *cysl-2* (a paralog of *cysl-1*) and the “O_2_-ON response” (38).

**Figure 2:**
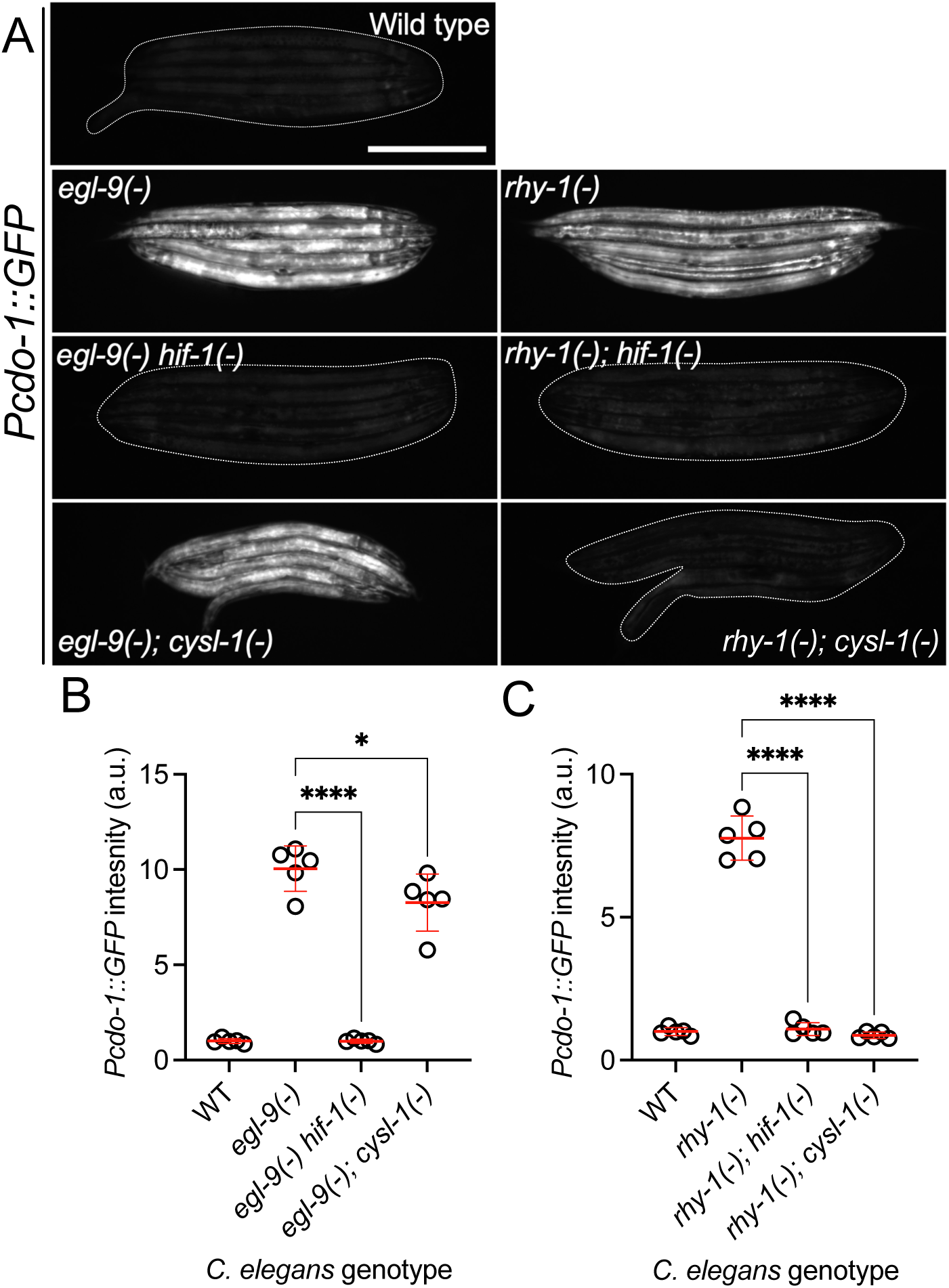
*cdo-1* transcription is activated by HIF-1 downstream of RHY-1, CYSL-1, and EGL-9. A) Expression of the *Pcdo-1::GFP* transgene is displayed for wild-type, *egl-9(sa307), egl-9(sa307) hif-1(ia4)* double mutant*, egl-9(sa307); cysl-1(ok762)* double mutant*, rhy-1(ok1402), rhy-1(ok1402); hif-1(ia4)* double mutant, and *rhy-1(ok1402); cysl-1(ok762)* double mutant *C. elegans* animals at the L4 stage of development. Scale bar is 250μm. White dotted line outlines animals with basal GFP expression. For GFP imaging, exposure time was 100ms. B,C) Quantification of the data displayed in Fig. 2A. Individual datapoints are shown (circles) as are the mean and standard deviation (red lines). *n* is 5 individuals per genotype. Data are normalized so that wild-type expression of *Pcdo-1::GFP* is 1 arbitrary unit (a.u.). *, p<0.05, ****, p<0.0001, ordinary one-way ANOVA with Dunnett’s post hoc analysis. Note, wild-type, *egl-9(-)*, and *rhy-1(-)* images in panel A and quantification of *Pcdo-1::GFP* in panels B and C are identical to the data presented in Fig. 1D,E. They are re-displayed here to allow for clear comparisons to the double mutant strains of interest.

To demonstrate that HIF-1 activates transcription of endogenous *cdo-1,* we explored published RNA-sequencing data of wild-type, *egl-9(-)*, and *egl-9(-) hif-1(-)* mutant animals (49). *egl-9(-)* mutant *C. elegans* display an 8-fold increase in *cdo-1* mRNA compared to wild type. This induction was dependent on *hif-1;* a *hif-1(-)* mutation completely suppressed the induction of *cdo-1* mRNA caused by an *egl-9(-)* mutation (49). These RNA-seq data confirm our findings using the *Pcdo-1::GFP* transcriptional reporter that HIF-1 is a transcriptional activator of *cdo-1.* ChIP-seq data of HIF-1 performed by the modERN project show that HIF-1 directly binds the *cdo-1* promoter (peak from −1,165 to −714 base pairs 5’ to the *cdo-1* ATG start codon) (50, 51). This HIF-1 binding site contains 3 copies of the HIF-binding motif (5’-RCGTG-3’) (52). Thus, *cdo-1* is a downstream effector of HIF-1 and is likely a direct transcriptional target of HIF-1.

### High levels of cysteine promote *cdo-1* transcription and cause lethality in *cysl-1* and *hif-1* mutant animals

Mammalian CDO1 levels and activity are highly induced by dietary cysteine (14–16). To determine if this homeostatic response is conserved in *C. elegans,* we exposed transgenic *C. elegans* carrying the *Pcdo-1::GFP* transcriptional reporter to high supplemental cysteine. Like our *egl-9* and *rhy-1* loss-of-function mutations, high levels of cysteine promoted *cdo-1* transcription (**Fig. 3A-C**). Despite the significant 3.6-fold induction of *Pcdo-1::GFP* caused by 100μM supplemental cysteine, we note that this induction is not as dramatic as the induction caused by null mutations in *egl-9* or *rhy-1.* This perhaps reflects the animals’ ability to buffer environmental cysteine which is bypassed by genetic intervention.

**Figure 3:**
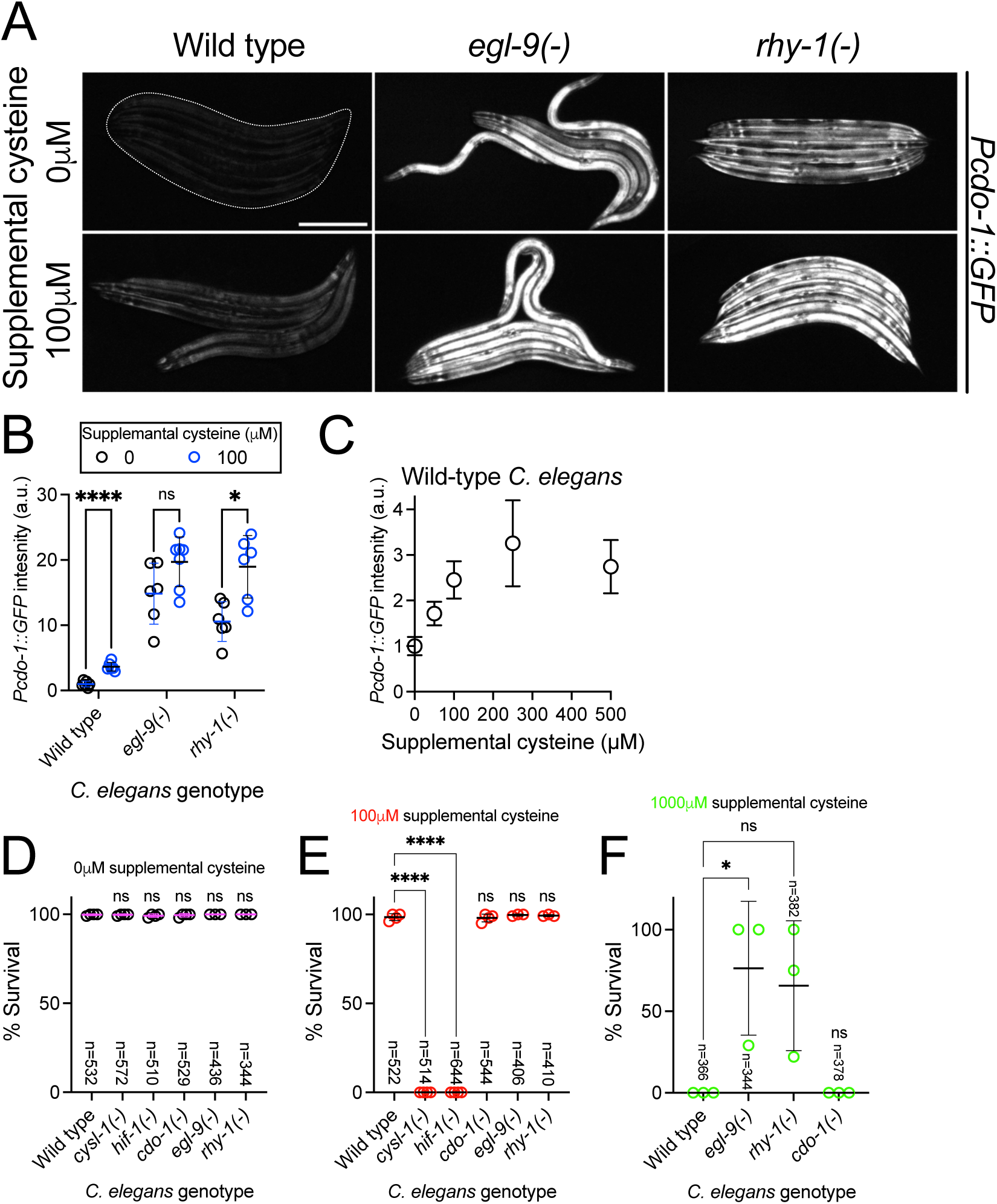
High levels of cysteine activate *cdo-1* transcription and are lethal to *hif-1* and *cysl-1* mutant animals. A) Expression of the *Pcdo-1::GFP* transgene is displayed for wild-type, *egl-9(sa307),* and *rhy-1(ok1402)* young-adult *C. elegans* exposed to 0 or 100μM supplemental cysteine. Scale bar is 250μm. White dotted line exposure time was 100ms. Supplemental cysteine did not impact the mortality of the animals being imaged. B) Quantification of the data displayed in Fig. 3A. Individual datapoints are shown (circles) as are the mean and standard deviation (black lines). *n* is 6 or 7 individuals per genotype. Data are normalized so that expression of *Pcdo-1::GFP* in wild-type *C. elegans* exposed to 0μM supplemental cysteine is equal to 1 arbitrary unit (a.u.). *, p<0.05, ****, p<0.0001, multiple unpaired t test with Welch’s correction. C) Quantification of the *Pcdo-1::GFP* expression is displayed for wild-type young-adult *C. elegans* exposed to 0, 50, 100, 250 or 500μM supplemental cysteine. Mean and standard deviation are displayed. n is 6 or 7 individuals per concentration of supplemental cysteine. D-F) The percentage of animals that survive overnight exposure to D) 0, E) 100, or F) 1,000μM supplemental cysteine. Individual datapoints (circles) represent biological replicates. 3 or 4 biological replicates were performed for each experiment and the total individuals scored amongst all replicates is displayed (n). *, p<0.05, ****, p<0.0001, ordinary one-way ANOVA with Dunnett’s post hoc analysis. ns indicates no significant difference was identified.

We hypothesized that cysteine might activate *cdo-1* transcription through the RHY-1/CYSL-1/EGL-9/HIF-1 pathway. In this pathway, *cysl-1* and *hif-1* act to promote *cdo-1* transcription. Thus, we sought to test whether *cysl-1* or *hif-1* were necessary for the induction of *Pcdo-1::GFP* by high levels of cysteine. However, this experiment was not possible as we observed 100% lethality in *cysl-1(-)* and *hif-1(-)* mutant animals exposed to 100μM supplemental cysteine, a cysteine concentration at which wild-type animals are healthy (**Fig. 3E**). Importantly, wild-type, *cysl-1(-)* and *hif-1(-)* animals were all healthy under control conditions without supplemental cysteine (**Fig. 3D**). While this phenotype limits our ability to establish the role of *cysl-1* and *hif-1* in the induction of *cdo-1* by high levels of cysteine, these data demonstrate that *cysl-1* and *hif-1* are necessary for survival under high cysteine conditions. Given the requirement of *hif-1* for survival in high levels of cysteine, we hypothesized that mutations in *egl-9* or *rhy-1* that activate *hif-1* might promote cysteine resistance. Indeed, we found that *egl-9(-)* and *rhy-1(-)* mutant *C. elegans* were partially viable when exposed to 1,000μM supplemental cysteine, a concentration that causes 100% lethality in wild-type animals (**Fig. 3F**). Thus, *egl-9* and *rhy-1* are negative regulators cysteine tolerance. Taken together, these data demonstrate a critical physiological role for the RHY-1/CYSL-1/EGL-9/HIF-1 pathway in promoting cysteine homeostasis.

To further test the interaction between high levels of cysteine and the RHY-1/CYSL-1/EGL-9/HIF-1 pathway, we exposed *egl-9(-); Pcdo-1::GFP* mutant animals to control or high levels of supplemental cysteine. We reasoned that if cysteine and *egl-9* loss of function promote *Pcdo-1::GFP* accumulation in the same pathway, then their effects should not be additive. Consistent with this hypothesis, we saw no difference in *Pcdo-1::GFP* expression in *egl-9(-)* mutant animals exposed to 0 or 100μM supplemental cysteine (**Fig. 3A,B**). We also tested the ability of supplemental cysteine to further activate *Pcdo-1::GFP* expression in *rhy-1(-)* mutant animals. In contrast to our results with *egl-9(-)*, we observed that high levels of cysteine caused a significant induction of *Pcdo-1::GFP* in the *rhy-1(-)* mutant background. These data suggest that cysteine acts in a pathway with *egl-9* but operates in parallel to the function of *rhy-1*.

Given the established role of CDO-1 in cysteine catabolism, we tested whether *cdo-1(-)* mutants were also sensitive to high levels of cysteine. *cdo-1(-)* mutant animals were not sensitive to high levels of supplemental cysteine compared to the wild type (**Fig. 3D-F**). Thus, *cdo-1* is not necessary for survival under high cysteine conditions. This suggests the existence of alternate pathways that promote cysteine homeostasis. Given the role of the RHY-1/CYSL-1/EGL-9/HIF-1 pathway in promoting cysteine homeostasis, we propose that HIF-1 activates pathways (in addition to *cdo-1*) that promote survival under high cysteine conditions.

### Activated CDO-1 accumulates and is functional in the hypodermis

We sought to identify the site of action of CDO-1. To observe CDO-1 localization, we used CRISPR/Cas9 to insert the GFP open reading frame into the endogenous *cdo-1* locus, replacing the native *cdo-1* stop codon. This *cdo-1(rae273)* allele encodes a C-terminal tagged “CDO-1::GFP” fusion protein from the native *cdo-1* genomic locus (**Fig. 4A**). To determine if CDO-1::GFP was functional, we combined the CDO-1::GFP fusion protein with a null mutation in *moc-1,* a gene that is essential for *C. elegans* Moco biosynthesis (33). We then observed the growth of *moc-1(-)* CDO-1::GFP *C. elegans* on wild-type and Moco-deficient *E. coli.* The *moc-1(-)* mutant *C. elegans* expressing CDO-1::GFP from the native *cdo-1* locus arrest during larval development when fed Moco-deficient *E. coli,* but not when fed Moco-producing *E.coli* (**Fig. 4C**). This lethality is caused by the CDO-1-mediated production of sulfites which are only toxic when *C. elegans* is Moco deficient, and demonstrates that the CDO-1::GFP fusion protein is functional (33). These data suggest that the CDO-1::GFP expression we observe is physiologically relevant.

**Figure 4:**
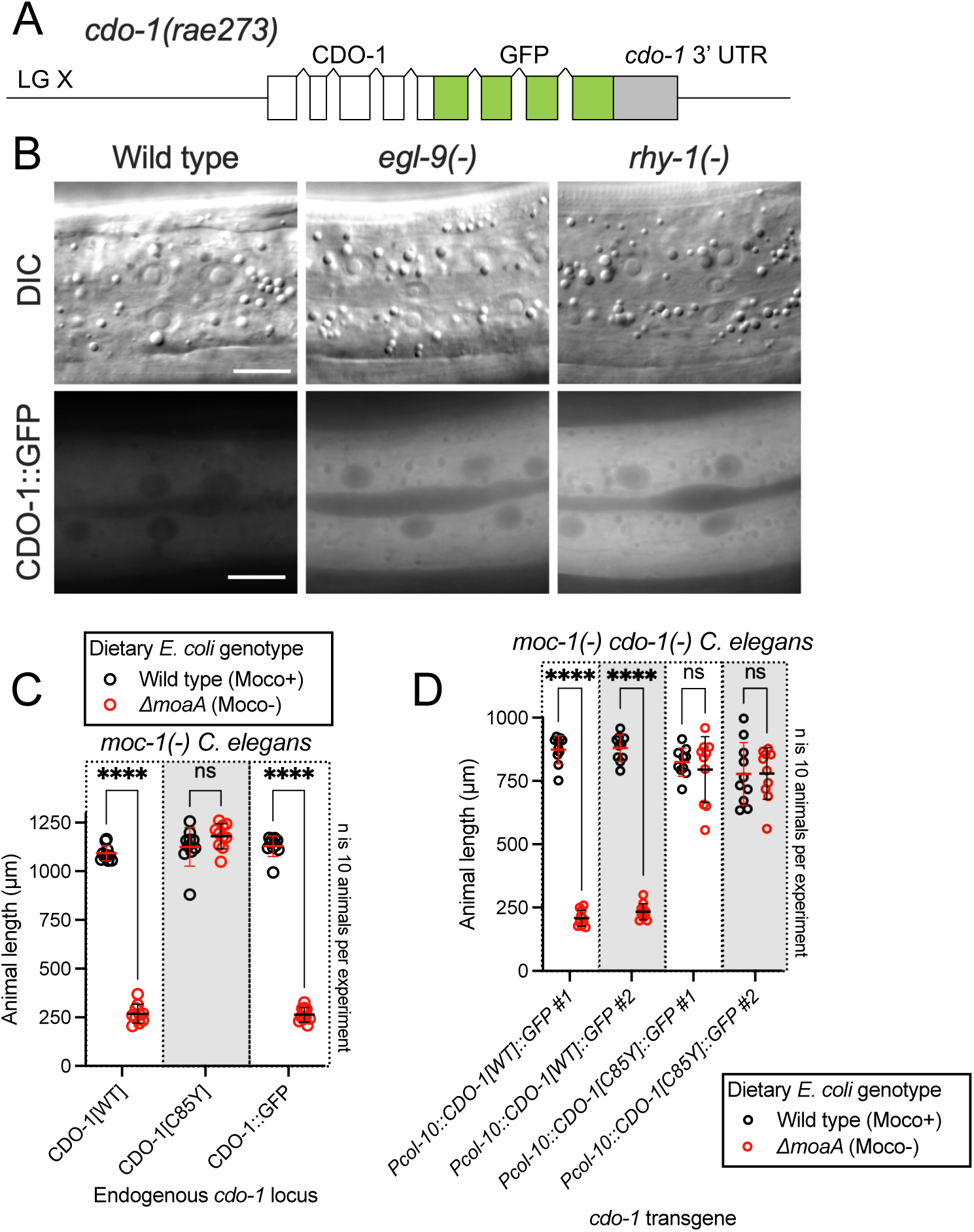
Hypodermal CDO-1 accumulates in the cytoplasm when *egl-9* or *rhy-1* are inactive and is sufficient to promote sulfur amino acid metabolism. A) Diagram of *cdo-1(rae273),* a CRISPR/Cas9-generated allele with GFP inserted into the *cdo-1* gene, creating a functional C-terminal CDO-1::GFP fusion protein expressed from the native *cdo-1* locus. B) Differential interference contrast (DIC) and fluorescence imaging are shown for wild-type, *egl-9(sa307),* and *rhy-1(ok1402) C. elegans* expressing CDO-1::GFP encoded by *cdo-1(rae273).* Scale bar is 10μm. For GFP imaging, exposure time was 200ms. An anterior segment of the Hyp7 hypodermal cell is displayed. CDO-1::GFP accumulates in the cytoplasm and is excluded from the nuclei. C) *moc-1(ok366), moc-1(ok366) cdo-1(mg622),* and *moc-1(ok366) cdo-1(rae273)* animals were cultured from synchronized L1 larvae for 72 hours on wild-type (black, Moco+) or Δ*moaA* mutant (red, Moco-) *E. coli.* D) *moc-1(ok366) cdo-1(mg622)* double mutant animals expressing *Pcol-10::CDO-1::GFP* or *Pcol-10::CDO-1[C85Y]::GFP* transgenes were cultured for 48 hours on wild-type (black, Moco+) or Δ*moaA* mutant (red, Moco-) *E. coli.* Two independently derived strains were tested for each transgene. For panels C and D, animal lengths were determined for each condition. Individual datapoints are shown (circles) as are the mean and standard deviation. Sample size (*n*) is 10 individuals for each experiment. ****, p<0.0001, multiple unpaired t test with Welch’s correction. ns indicates no significant difference was identified.

When wild-type animals expressing CDO-1::GFP were grown under standard culture conditions, we observed CDO-1::GFP expression in multiple tissues, including prominent expression in the hypodermis (**Fig. 4B, Fig. S2**). We tested if CDO-1::GFP fusion protein levels were affected by inactivation of *egl-9* or *rhy-1.* We generated *egl-9(-)*; CDO-1::GFP and *rhy-1(-)*; CDO-1::GFP animals, and assayed expression of CDO-1::GFP. We found that CDO-1::GFP fusion protein levels, encoded by *cdo-1(rae273),* were increased by *egl-9(-)* or *rhy-1(-)* mutations (**Fig. 4B, Fig. S2**). This is consistent with our studies using *Pcdo-1::CDO-1::GFP* and *Pcdo-1::GFP* transgenes. Furthermore, high levels of cysteine also promoted accumulation of CDO-1::GFP (**Fig. S3**). In all scenarios, the hypodermis was the most prominent site of CDO-1::GFP accumulation. Specifically, we found that CDO-1::GFP was expressed in the cytoplasm of Hyp7, the major *C. elegans* hypodermal cell (**Fig. 4B**).

Based on the expression pattern of CDO-1::GFP encoded by *cdo-1(rae273)*, we hypothesized that CDO-1 acts in the hypodermis to promote sulfur amino acid metabolism. To test this hypothesis, we engineered a *cdo-1* rescue construct in which *cdo-1* is expressed exclusively in the hypodermis under the control of a hypoderm-specific collagen (*col-10*) promoter (*Pcol-10::CDO-1::GFP*) (53). Collagens are expressed exclusively in the hypodermis of nematodes, with a periodic induction in phase with the almost diurnal molting cycle (54). Multiple independent transgenic *C. elegans* strains were generated by integrating the *Pcol-10::CDO-1::GFP* construct into the *C. elegans* genome and tested for rescue of the *cdo-1(-)* mutant suppression of Moco-deficient larval arrest (41). Thus, tissue-specific complementation of the *cdo-1* mutation would regenerate a lethal arrest phenotype caused by Moco deficiency. We found that multiple independent transgenic strains of *cdo-1(-) moc-1(-)* double mutant animals carrying the *Pcol-10::CDO-1::GFP* transgene displayed a larval arrest phenotype when fed a Moco-deficient diet (**Fig. 4D**). These data demonstrated that hypodermal-specific expression of *cdo-1* is sufficient to rescue the *cdo-1(-)* mutant suppression of Moco-deficient lethality. This rescue was dependent upon the enzymatic activity of CDO-1 as an active site variant of this transgene (*Pcol-10::CDO-1[C85Y]::GFP)* did not rescue the suppressed larval arrest of *cdo-1(-) moc-1(-)* double mutant animals fed Moco-deficient diets (**Fig. 4D**). Taken together, our analyses of CDO-1::GFP expression demonstrate that CDO-1 is expressed, and that expression is regulated, in multiple tissues, principal among them being the hypodermis and Hyp7 cell. Our tissue-specific rescue data demonstrate that hypodermal expression of *cdo-1* is sufficient to promote cysteine catabolism and suggest that the hypodermis is a critical tissue for sulfur amino acid metabolism. However, we cannot exclude the possibility that CDO-1 also acts in other cells and tissues as well.

### HIF-1 promotes CDO-1 activity downstream of the H_2_S-sensing pathway

We sought to determine the physiological impact of *cdo-1* transcriptional activation by HIF-1. We reasoned mutations that activate HIF-1 and increase *cdo-1* transcription may cause increased CDO-1 activity. CDO-1 sits at a critical metabolic node in the degradation of the sulfur amino acids cysteine and methionine (**Fig 1A**). A key byproduct of sulfur amino acid metabolism and CDO-1 is sulfite, a reactive toxin that is detoxified by the Moco-requiring sulfite oxidase (SUOX-1). Null mutations in *suox-1* cause larval lethality. However, animals carrying the *suox-1(gk738847)* hypomorphic allele are healthy under standard culture conditions (33, 34). *suox-1(gk738847)* mutant animals display only 4% SUOX-1 activity compared to wild type and are exquisitely sensitive to sulfite stress (55). Thus, the *suox-1(gk738847)* mutation creates a sensitized genetic background to probe for increases in endogenous sulfite production. To test if increased *cdo-1* transcription would impact the growth of *suox-1-*comprimised animals, we combined the *egl-9* null mutation, which promotes HIF-1 activity and *cdo-1* transcription, with the *suox-1(gk738847)* allele. While *egl-9(-)* and *suox-1(gk738847)* single mutant animals are healthy under standard culture conditions, the *egl-9(-); suox-1(gk738847)* double mutant animals are extremely sick and require significantly more days to exhaust their *E. coli* food source under standard culture conditions (**Table 1**). These data establish a synthetic genetic interaction between these loci. To determine the role of sulfur amino acid metabolism in the *egl-9(-); suox-1(gk738847)* synthetic sickness phenotype, we engineered *egl-9(-); cdo-1(-) suox-1(gk738847)* and *cth-2(-); egl-9(-); suox-1(gk738847)* triple mutant animals. The *egl-9; suox-1* synthetic sickness phenotype was suppressed by inactivating mutations in *cdo-1* or *cth-2* which block the endogenous production of sulfite (**Table 1**). These data demonstrate that the deleterious activity of the *egl-9(-)* mutation in a *suox-1(gk738847)* background requires functional sulfur amino acid metabolism.

**Table 1:** Growth of *C. elegans* strains on standard laboratory conditions. *C. elegans* strains and their corresponding mutations are displayed. For each strain, 5 L4-stage animals were seeded onto standard NGM petri dishes seeded with a monoculture of *E. coli* OP50. Petri dishes were monitored until the animals (and their progeny) depleted the lawn of *E. coli*. This was recorded as “days for population to starve petri dish”. The average of these experiments is displayed for each *C. elegans* strain as is the standard deviation (SD) and the number of biological replicates (*n*). Significant differences in population growth were determined by appropriate comparisons to either *suox-1(gk738847)* (GR2269) or *egl-9(sa307); suox-1(gk738847)* (USD421) using an ordinary one-way ANOVA with Dunnett’s post hoc analysis. No test indicates a statistical comparison was not made. Note, the data for USD421 are displayed twice in the table to allow for ease of comparison.

| *C. elegans* strain | Genetic locus 1 | Genetic locus 2 | Genetic locus 3 | Days for population to starve<br>petri dish ± SD (n) | Significant difference<br>compared to GR2269<br>(Adjusted P Value) |
| --- | --- | --- | --- | --- | --- |
| N2 | Wild type | Wild type | Wild type | 5±1 (8) | No test |
| GR2269 | Wild type | *suox-1(gk738847)* | Wild type | 8±1 (4) | No test |
| JT307 | *egl-9(sa307)* | Wild type | Wild type | 7±1 (5) | No test |
| USD421 | *egl-9(sa307)* | *suox-1(gk738847)* | Wild type | 19±5 (11) | 0.0003 (***) |
| USD926 | *egl-9(rae276)* | Wild type | Wild type | 6±1 (3) | No test |
| USD937 | *egl-9(rae276)* | *suox-1(gk738847)* | Wild type | 9±1 (3) | 0.93 (ns) |
| USD512 | *rhy-1(ok1402)* | Wild type | Wild type | 6±0 (4) | No test |
| USD414 | *rhy-1(ok1402)* | *suox-1(gk738847)* | Wild type | 7±0 (4) | 0.99 (ns) |
| CB5602 | *vhl-1(ok161)* | Wild type | Wild type | 6±0 (4) | No test |
| USD422 | *vhl-1(ok161)* | *suox-1(gk738847)* | Wild type | 8±1 (4) | 0.99 (ns) |
| *C. elegans* strain | Genetic locus 1 | Genetic locus 2 | Genetic locus 3 | Days for population to starve<br>petri dish ± SD (n) | Significant difference<br>compared to USD421<br>(Adjusted P value) |
| USD421 | *egl-9(sa307)* | *suox-1(gk738847)* | Wild type | 19±5 (11) | No test |
| USD430 | *egl-9(sa307)* | *suox-1(gk738847)* | *cdo-1(mg622)* | 7±0 (6) | <0.0001 (****) |
| USD433 | *egl-9(sa307)* | *suox-1(gk738847)* | *cth-2(mg599)* | 9±1 (4) | <0.0001 (****) |
| USD432 | *egl-9(sa307)* | *suox-1(gk738847)* | *rhy-1(ok1402)* | 7±1 (4) | <0.0001 (****) |
| USD434 | *egl-9(sa307)* | *suox-1(gk738847)* | *cysl-1(ok762)* | 16±1 (9) | 0.1 (ns) |
| USD431 | *egl-9(sa307)* | *suox-1(gk738847)* | *hif-1(ia4)* | 9±1 (6) | <0.0001 (****) |

To determine the role of the H_2_S-sensing pathway in the synthetic sickness phenotype displayed by *egl-9(-); suox-1(gk738847)* double mutant animals, we introduced null alleles of *cysl-1* or *hif-1* into the *egl-9(-); suox-1(gk738847)* double mutant. The *egl-9; suox-1* synthetic sickness phenotype was dependent upon *hif-1* but not *cysl-1* (**Table 1**). These results are consistent with our proposed genetic pathway and support the model that transcriptional activation of *cdo-1* by HIF-1 causes increased CDO-1 activity and increased flux of sulfur amino acids through their catabolic pathway.

Loss of *rhy-1* also strongly activates *cdo-1* transcription. We hypothesized that *rhy-1(-); suox-1(gk738847)* double mutant animals would display a synthetic sickness phenotype like *egl-9(-); suox-1(gk738847)* double mutant animals. However, based upon their ability to exhaust their *E. coli* food source under standard culture conditions, *rhy-1(-); suox-1(gk738847)* double mutant animals were just as healthy as either *rhy-1(-)* or *suox-1(gk738847)* single mutant *C. elegans* (**Table 1**). These data suggest that increasing *cdo-1* transcription alone is not sufficient to promote sulfite production via CDO-1. In addition to the role played by *rhy-1* in the regulation of HIF-1 activity, *rhy-1* itself is a transcriptional target of HIF-1. Loss of *egl-9* activity induces *rhy-1* mRNA ∼50-fold in a *hif-1-*dependent manner (49). Given this potent transcriptional activation, we wondered if *rhy-1* might play an additional role downstream of HIF-1 in the regulation of sulfur amino acid metabolism and sulfite production. To test this hypothesis, we engineered *rhy-1(-); egl-9(-); suox-1(gk738847)* triple mutant animals. To our surprise, the *rhy-1(-); egl-9(-); suox-1(gk738847)* triple mutant animals were healthy, demonstrating that *rhy-1* was necessary for the deleterious activity of the *egl-9(-)* mutation in a *suox-1(gk738847)* background (**Table 1**). These genetic data suggest a dual role for *rhy-1* in the control of sulfur amino acid metabolism; first as a component of a regulatory cascade that controls the activity of HIF-1 and second as a functional downstream effector of HIF-1 that is required for sulfur amino acid metabolism.

This is not the first description of a *rhy-1* role downstream of *hif-1.* Overexpression of a *rhy-1*-encoding transgene suppresses the lethality of a *hif-1(-)* mutant during H_S_S stress (56). These data establish RHY-1 as both a regulator and effector of HIF-1. How RHY-1, a predicted membrane-bound O-acyltransferase, molecularly executes these dual roles remains to be explored.

### EGL-9 prolyl hydroxylase activity and VHL-1 are largely dispensable in the regulation of CDO-1

EGL-9 inhibits HIF-1 through its prolyl hydroxylase domain that hydroxylates HIF-1 proline 621 (**Fig. 5A**) (39). To evaluate the impact of the EGL-9 prolyl hydroxylase domain on the regulation of *cdo-1*, we generated a prolyl hydroxylase domain-inactive *egl-9* mutation using CRISPR/Cas9. We engineered an *egl-9* mutation that substitutes an alanine in place of histidine 487 (H487A) (**Fig. 1C**). Histidine 487 of EGL-9 is highly conserved and catalytically essential in the prolyl hydroxylase domain active site (**Fig. 5B**) (57, 58). To evaluate the impact of an inactive EGL-9 prolyl hydroxylase domain on the transcription of *cdo-1,* we engineered a *C. elegans* strain carrying the *egl-9(H487A)* mutation with the *Pcdo-1::GFP* transcriptional reporter. *egl-9(H487A)* caused a modest increase in *Pcdo-1::GFP* accumulation in the *C. elegans* intestine, suggesting that the EGL-9 prolyl hydroxylase domain is necessary to repress *cdo-1* transcription in the intestine (**Fig. 5C,D**). However, the activation of *Pcdo-1::GFP* by the *egl-9(H487A)* mutation was markedly less when compared to *Pcdo-1::GFP* activation caused by an *egl-9(-)* null mutation (**Fig. 5C,D**). These data suggest that EGL-9 has a prolyl hydroxylase domain-independent activity that is responsible for repressing *cdo-1* transcription.

**Figure 5:**
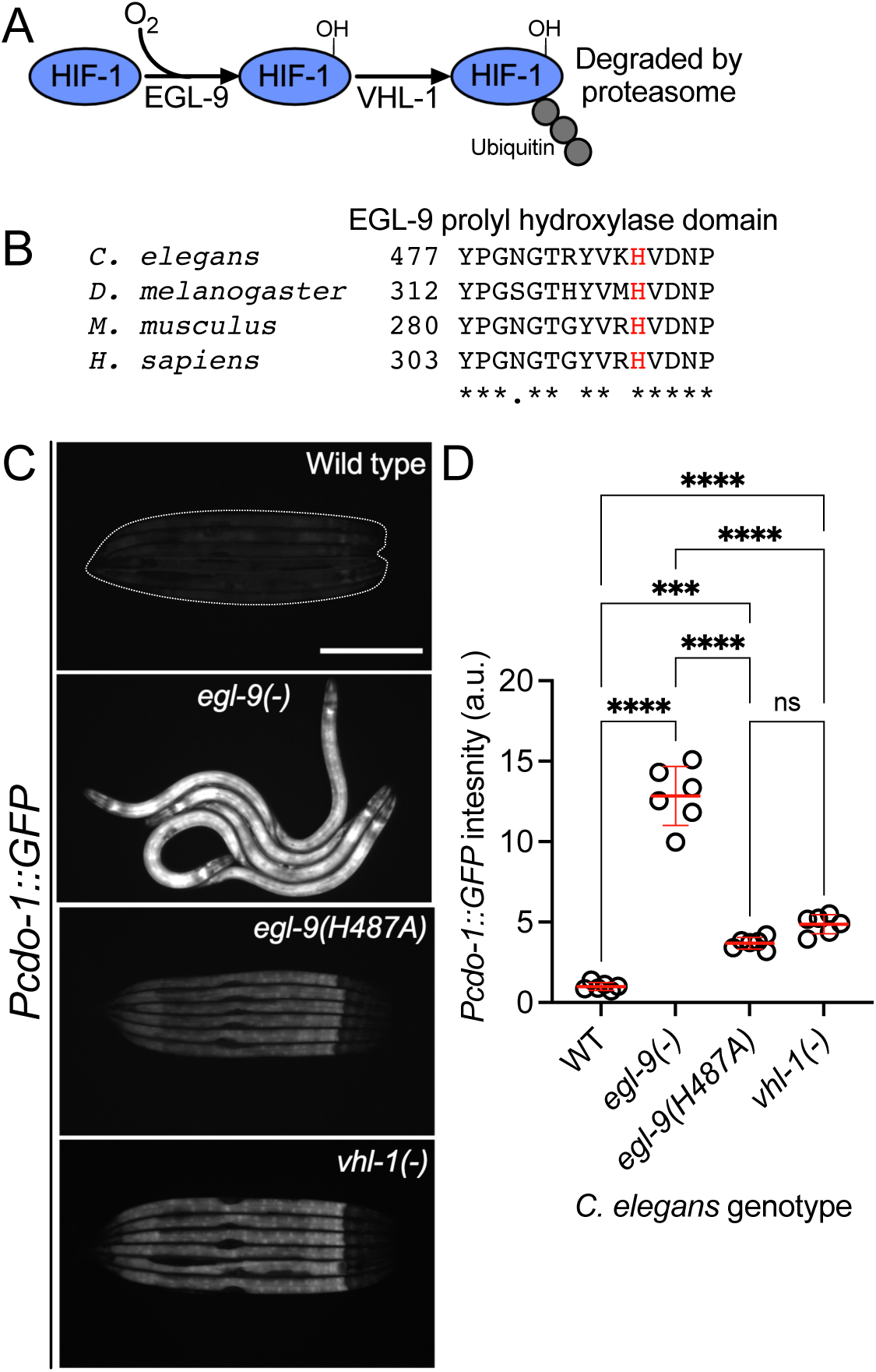
*egl-9* inhibits *cdo-1* transcription in a largely prolyl-hydroxylase and VHL-1-independent manner. A) The pathway of HIF-1 processing during normoxia is displayed. EGL-9 uses O_2_ as a substrate to hydroxylate (-OH) HIF-1 on specific proline residues. Prolyl hydroxylated HIF-1 is bound by VHL-1 which facilitates HIF-1 polyubiquitination and targets HIF-1 for degradation by the proteasome. B) Amino acid alignment of the EGL-9 prolyl hydroxylase domain from *C. elegans, D. melanogaster, M. musculus,* and *H. sapiens.* “*” indicate perfect amino acid conservation while “.” indicates weak similarity amongst species compared. Highlighted (red) is the catalytically essential histidine 487 residue in *C. elegans*. Alignment was performed using Clustal Omega (EMBL-EBI). C) Expression of *Pcdo-1::GFP* promoter fusion transgene is displayed for wild-type, *egl-9(sa307, -), egl-9(rae276,* H487A), and *vhl-1(ok161) C. elegans* animals at the L4 stage of development. Scale bar is 250μm. White dotted line outlines animals with basal GFP expression. For GFP imaging, exposure time was 500ms. D) Quantification of the data displayed in Fig. 5C. Individual datapoints are shown (circles) as are the mean and standard deviation (red lines). *n* is 6 individuals per genotype. Data are normalized so that wild-type expression of *Pcdo-1::GFP* is 1 arbitrary unit (a.u.). ***, p<0.001, ****, p<0.0001, ordinary one-way ANOVA with Tukey’s multiple comparisons test. ns indicates no significant difference was identified.

EGL-9 hydroxylates specific proline residues on HIF-1. These hydroxylated proline residues are recognized by the von Hippel-Lindau E3 ubiquitin ligase (VHL-1). VHL-1-mediated ubiquitination promotes degradation of HIF-1 by the proteasome (**Fig. 5A**) (40). Thus, the EGL-9 prolyl hydroxylase domain and VHL-1 act in a pathway to regulate HIF-1. To determine the role of VHL-1 in regulating *cdo-1,* we engineered a *C. elegans* strain carrying the *vhl-1(-)* null mutation with our P*cdo-1::GFP* reporter. *vhl-1* inactivation also caused a modest increase in *Pcdo-1::GFP* expression in the *C. elegans* intestine, suggesting that *vhl-1* is necessary to repress *cdo-1* transcription (**Fig. 5C,D**). However, activation of *Pcdo-1::GFP* by the *vhl-1(-)* mutation was less than activation caused by an *egl-9(-)* null mutation (**Fig. 5C,D**). These data suggest that EGL-9 has a VHL-1-independent activity that is responsible for repressing *cdo-1* transcription.

To evaluate the impact of inactivating the EGL-9 prolyl hydroxylase domain or VHL-1 on cysteine metabolism, we again employed the *suox-1(gk738847)* hypomorphic mutation that sensitizes animals to increases in sulfite. We engineered *egl-9(H487A); suox-1(gk738847)* and *vhl-1(-) suox-1(gk738847)* double mutant animals and evaluated the health of those strains. In contrast to *egl-9(-); suox-1(gk738847)* double mutant animals which are extremely sick, *egl-9(H487A); suox-1(gk738847)* and *vhl-1(-) suox-1(gk738847)* double mutant animals are healthy (**Table 1**). These genetic data suggest that neither the EGL-9 prolyl hydroxylase domain nor VHL-1 are necessary to repress cysteine catabolism/sulfite production. However, it is plausible that the *egl-9(H487)* or *vhl-1(-)* mutations modestly activate cysteine metabolism, likely proportional to their activation of the *Pcdo-1::GFP* transgene (**Fig. 5C,D**), and that this activation is not sufficient to produce enough sulfites to negatively impact the growth of *suox-1(gk738847)* mutant animals.

## Discussion

### CDO-1 is a physiologically relevant effector of HIF-1

The hypoxia-inducible factor HIF-1 is a master regulator of the cellular response to hypoxia. It activates the transcription of many genes and pathways that are critical to maintain metabolic homeostasis in the face of low O_2_. For example, mammalian HIF1α induces the hematopoietic growth hormone erythropoietin, glucose transport and glycolysis, as well as lactate dehydrogenase (59–62). The nematode *C. elegans* encounters a range of O_2_ tensions in its natural habitat of rotting material: as microbial abundance increases, O_2_ levels decrease from atmospheric levels (∼21% O_2_). In fact, *C. elegans* prefers 5-12% O_2_, perhaps because hypoxia predicts abundant bacterial food sources (63). Members of the HIF-1 pathway and its targets have emerged from genetic studies of *C. elegans* (39). For instance, *C. elegans* HIF-1 promotes H_2_S homeostasis by inducing transcription of the mitochondrial sulfide quinone oxidoreductase (*sqrd-1),* detoxifying H_2_S (37).

We sought to define genes that regulate the levels and activity of cysteine dioxygenase (CDO-1), a critical regulator of cysteine homeostasis (14–16). Taking an unbiased genetic approach in *C. elegans,* we found that *cdo-1* was highly regulated by HIF-1 downstream of a signaling pathway that includes *rhy-1*, *cysl-1*, and *egl-9*. We demonstrated that HIF-1 promotes *cdo-1* transcription, accumulation of CDO-1 protein, and increased CDO-1 activity. Loss of *rhy-1* or *egl-9* activate *hif-1* to in turn strongly induce the *cdo-1* promoter. The pathway for activation of *cdo-1* also requires *cysl-1,* which functions downstream of *rhy-1* and upstream of *egl-9.* Based on ChIP-Seq studies, HIF-1 directly binds to the *cdo-1* promoter (50). Interestingly, mammalian CDO1 is regulated at both the transcriptional and post-transcriptional level in response to high dietary cysteine (17–21). Our studies in the nematode *C. elegans* suggest that CDO1 transcription in mammals might be governed by HIF1α in response to changes in cellular cysteine. Importantly, all members of the RHY-1/CYSL-1/EGL-9 pathway in *C. elegans* have homologs encoded by mammalian genomes (38). Given the conservation of these proteins, future studies may show that similar cysteine and H_2_S-responsive signaling pathways operate in mammals.

We also show that the transcriptional activation of *cdo-1* by HIF-1 promotes CDO-1 enzymatic activity. Genetic activation of *cdo-1* by loss of *egl-9* causes severe sickness in a mutant with reduced sulfite oxidase activity, an activity required to cope with the toxic sulfite produced via CDO-1. This synthetic sickness is dependent upon an intact HIF-1 signaling pathway and a functioning sulfur amino acid metabolism pathway, as the dramatic sickness of an *egl-9; suox-1* double mutant is suppressed by loss of *hif-1, cth-2,* or *cdo-1.* These genetic results validate the intersection of HIF-1 and CDO-1 in converging biological regulatory pathways and encourage further exploration of this regulatory node. The potential physiological relevance of the connection between HIF-1 signaling and cysteine metabolism is discussed below.

It was initially surprising that the canonical hypoxia-sensing transcription factor, that is activated by relatively low O_2_ tensions, induces transcription of *cdo-1,* which encodes an oxidase that requires dissolved O_2_ to oxidize cysteine. Both EGL-9 and CDO-1 are dioxygenases, requiring O_2_ as a substrate to catalyze their chemical reactions. EGL-9 functions as an O_2_ sensor, using its O_2_ substrate to hydroxylate HIF-1 at relatively high O_2_ partial pressures, targeting it for degradation. EGL-9 is uniquely poised to sense deviations from the normally high physiological O_2_ concentrations given its high K_m_ for O_2_ (64–68). While it is not known if mammalian CDO1 or *C. elegans* CDO-1 dioxygenases are active at lower O_2_ tensions than EGL-9, it is possible, even probable, that the set point for activation of HIF-1 by the failure of EGL-9 prolyl hydroxylation, is at a higher O_2_ tension than the K_m_ of CDO-1 for oxidation of cysteine. In this way, CDO-1 could still coordinate cysteine catabolism while the cell is experiencing hypoxia and EGL-9 is unable to hydroxylate HIF-1.

### Evidence for distinct pathways of HIF-1 activation by hypoxia and cysteine/H_2_S

The hypoxia-signaling pathway is defined by EGL-9-dependent prolyl hydroxylation of HIF-1. Hydroxylated HIF-1 is then targeted for degradation via VHL-1-mediated ubiquitination (39, 40, 52). However, multiple lines of evidence, reinforced by our work, demonstrate VHL-1- and prolyl hydroxylase-independent activity of EGL-9. *egl-9(-)* null mutant *C. elegans* accumulate HIF-1 protein and display increased transcription of many genes, including *nhr-57,* an established target of HIF-1 (36). In rescue experiments of an *egl-9(-)* null mutant, Shao *et al.* (2009) demonstrate that a wild-type *egl-9* transgene restores normal HIF-1 protein levels and HIF-1 transcription. However, rescue experiments with a prolyl hydroxylase domain-inactive *egl-9(H487A)* transgene do not correct the accumulation of HIF-1 protein and only partially reduce the HIF-1 transcriptional output (58). Through our studies of *cdo-1* transcription, we demonstrate that an *egl-9(H487A)* mutant incompletely activates HIF-1 transcription when compared to an *egl-9(-)* null mutation (**Fig. 5C,D).** Taken together, we conclude EGL-9 has activity independent of its prolyl hydroxylase domain, mirroring and supporting previous work (58).

In studies of the HIF-1-dependent P*nhr-57::GFP* transcriptional reporter, Shen *et al.* (2006) observed that *egl-9(-)* null mutants promote HIF-1 transcription more than a *vhl-1(-)* mutant (36). We observe this same distinction between *egl-9* and *vhl-1* mutations with our *Pcdo-1::GFP* reporter (**Fig. 5C,D).** This difference in transcriptional activity is not explained by HIF-1 protein levels as HIF-1 protein accumulated equally in *egl-9(-)* and *vhl-1(-)* null mutants (36). This observation suggests that EGL-9 represses both HIF-1 levels and activity. This study also notes that *vhl-1* represses HIF-1 transcription in the intestine while *egl-9* acts in a wider array of tissues including the intestine and the hypodermis (36). Budde and Roth (2010) also demonstrate that loss of *vhl-1* promotes HIF-1 transcription in the *C. elegans* intestine while total loss of *egl-9* promotes HIF-1 transcription in multiple tissues including the intestine and the hypodermis (69). Our data expand upon these observations by demonstrating that loss of the EGL-9 prolyl hydroxylase domain promotes *cdo-1* transcription in the intestine alone, mirroring the loss of *vhl-1* **(Fig. 5C**). Budde and Roth (2010) additionally demonstrate that physiologically relevant stimuli also elicit a tissue-specific transcriptional response: hypoxia promotes HIF-1 transcription in the intestine while H_2_S promotes HIF-1 transcription in the hypodermis. Importantly, H_2_S promotes hypodermal HIF-1 transcription in a *vhl-1(-)* mutant, demonstrating a VHL-1-independent pathway for H_2_S activation of HIF-1 (69). We demonstrate that high levels of cysteine promote *cdo-1* transcription in the hypodermis. Taken together with our data, these studies suggest two distinct pathways for activating HIF-1 transcription: i) a hypoxia-sensing pathway that is dependent upon *vhl-1* and the EGL-9 prolyl hydroxylase domain and promotes HIF-1 activity in the intestine and ii) an H_2_S-sensing pathway that is independent of *vhl-1* and the EGL-9 prolyl hydroxylase domain and promotes HIF-1 activity in the hypodermis.

The focus of CDO-1 expression and regulation of sulfite production in the hypoderm may be due to the demand in sulfur metabolism as well as oxygen-dependent hydroxylation of collagens during the *C. elegans* molting cycle. Many collagen genes are expressed before each larval molt and many collagen prolines are hydroxylated, like particular HIF-1 prolines, and their cysteines form disulfides during collagen assembly. The large demand for the amino acid cysteine in the many collagen genes expressed at high levels during a molt may challenge cysteine homeostasis in the hypodermis (54).

The genetic details of the H_2_S-sensing pathway were solidified through studies of *rhy-1* in *C. elegans.* Ma *et al.* (2012) demonstrate that *hif-1* repression via *rhy-1* requires the activity of *cysl-1,* a gene encoding a cysteine synthase-like protein (38). They further demonstrate that high H_2_S promotes a physical interaction between CYSL-1 and EGL-9, resulting in the inactivation of EGL-9 and increased HIF-1 activity. Together, these studies suggest distinct genetic regulators of EGL-9/HIF-1 signaling: *rhy-1* and *cysl-1* govern the H_2_S-sensing pathway while *vhl-1* mediates the hypoxia-sensing pathway. These pathways are distinct in their requirement for the EGL-9 prolyl hydroxylase domain.

### A negative feedback loop senses high cysteine/H_2_S, promotes CDO-1 activity, and maintains cysteine homeostasis

Why would the RHY-1/CYSL-1/EGL-9/HIF-1 H_2_S-sensing pathway control the levels and activity of cysteine dioxygenase? We speculate this intersection facilitates a homeostatic pathway allowing *C. elegans* to sense and respond to cysteine level. We propose that H_2_S acts as a gaseous signaling molecule to promote cysteine catabolism. H_2_S activates HIF-1 in the hypodermis by promoting the CYSL-1-mediated inactivation of EGL-9 (38). We show that high levels of cysteine similarly induce *cdo-1* transcription in the hypodermis. Our genetic data demonstrate that *cdo-1* is induced by the same genetic pathway that senses H_2_S in *C. elegans* and CDO-1 acts in the hypodermis, the major site of H_2_S-induced transcription. Furthermore, H_2_S induces endogenous *cdo-1* transcription >3-fold while *cdo-1* mRNA levels do not change when *C. elegans* are exposed to hypoxia (70, 71). Thus, it is likely that H_2_S promotes *cdo-1* transcription through RHY-1, CYSL-1, EGL-9, and HIF-1. H_2_S is a reasonable small molecule signal to alert cells to high levels of cysteine. Excess cysteine results in the production of H_2_S mediated by multiple enzymes including cystathionase (CTH), cystathionine β-synthase (CBS), and 3-mercaptopyruvate sulfurtransferase (MST) (9, 72). For example, CTH activity within the carotid body produces H_2_S that modifies the mammalian response to hypoxia (73). We speculate that excess cysteine in *C. elegans* promotes the enzymatic production of H_2_S which activates HIF-1 via the RHY-1/CYSL-1/EGL-9 signaling pathway. In our homeostatic model, H_2_S-activated HIF-1 would then induce *cdo-1* transcription, promoting CDO-1 activity and the catabolism of the high-cysteine trigger (**Fig. 6**). Supporting this model, *cysl-1(-)* and *hif-1(-)* mutant *C. elegans* cannot survive in a high cysteine environment, demonstrating their central role in promoting cysteine homeostasis.

**Figure 6:**
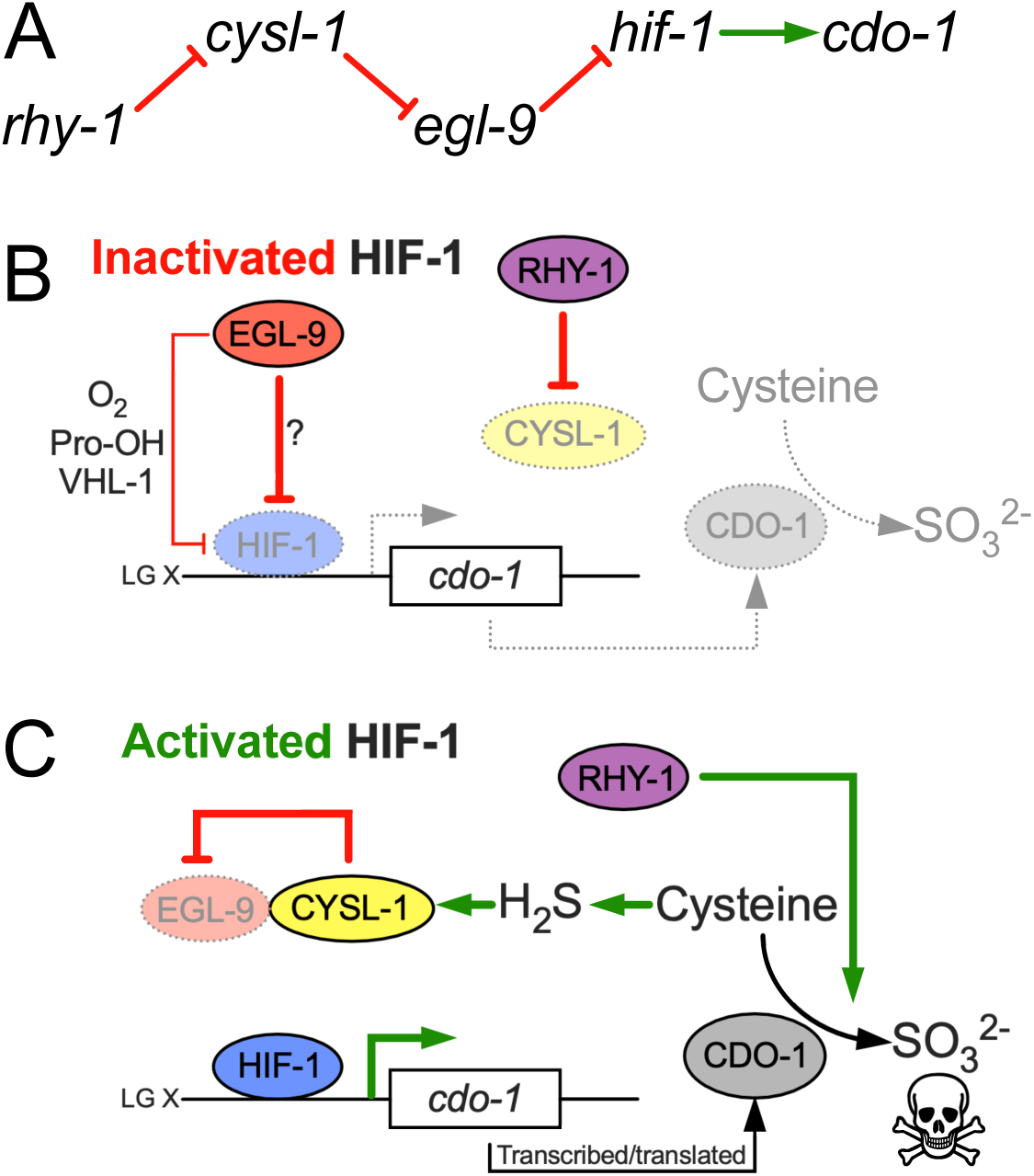
Model for the regulation of cysteine metabolism by HIF-1. A) Proposed genetic pathway for the regulation of *cdo-1. rhy-1, cysl-1,* and *egl-9* act in a negative-regulatory cascade to control activity of the HIF-1 transcription factor, which activates transcription of *cdo-1.* B) Under basal conditions (inactivated HIF-1), EGL-9 negatively regulates HIF-1 through 2 distinct pathways; one pathway is dependent upon O_2_, prolyl hydroxylation (Pro-OH), and VHL-1, while the second acts independently of these canonical factors. Under these conditions *cdo-1* transcription is kept at basal levels and cysteine catabolism is not induced. C) During conditions where HIF-1 is activated (high H_2_S), CYSL-1 directly binds and inhibits EGL-9, preventing HIF-1 inactivation. Active HIF-1 binds the *cdo-1* promoter, driving transcription and promoting CDO-1 protein accumulation. High CDO-1 levels promote the catabolism of cysteine leading to production of sulfites (SO_3_^2-^) that are toxic during Moco or SUOX-1 deficiency. HIF-1-induced cysteine catabolism requires the activity of *rhy-1*.

Importantly, the human ortholog of EGL-9 (EglN1) has previously been implicated as a cysteine sensor in the context of triple negative breast cancer (TNBC) (74). This study determined that HIF1α accumulates in TNBC cells even during normoxia. Briggs *et al.* (2016) demonstrate that L-glutamate secretion via TNBC cells suppresses HIF1α prolyl hydroxylation, stabilizing HIF1α. L-glutamate secretion is mediated via the glutamate/cystine antiporter, xCT (75, 76). L-glutamate secretion inhibits xCT and a concomitant decrease in intracellular cystine/cysteine was observed. The authors propose that low intracellular cystine/cysteine produces oxidizing conditions that oxidize specific EglN1 cysteine residues, inhibiting the activity of EglN1. Thus, Briggs *et al.* (2016) propose EglN1 as a cysteine sensor whose activity is promoted by cysteine.

Our work in *C. elegans* also strongly suggests a role for EGL-9 in sensing and responding to cysteine. We show that high levels of cysteine promote transcription of the HIF-1 target gene *cdo-1,* and that this induction is not additive with a null mutation in *egl-9,* suggesting cysteine and *egl-9* act in a pathway. Therefore, we propose that in *C. elegans* high levels of cysteine inhibit the activity of EGL-9, the opposite effect observed in TNBC cells. Furthermore, our genetic studies demonstrate that regulation of *cdo-1* transcription occurs largely independent of the EGL-9 prolyl hydroxylase domain and VHL-1, while the mechanism proposed by Briggs *et al.* (2016) suggests that the stabilization of HIF1α by L-glutamate is correlated with increased prolyl hydroxylation. Despite these differences, it seems likely that the roles for *C. elegans* EGL-9 and human EglN1 in sensing cysteine are connected. However, additional studies are required to determine if there is an evolutionary relationship between these cysteine-sensing mechanisms.

Members of the H_2_S-sensing pathway have also been implicated in the *C. elegans* response to infection (44, 77). Many secreted proteins that mediate intercellular signaling or innate immune responses to pathogens are cysteine-rich and form disulfide bonds in the endoplasmic reticulum before protein secretion (78, 79). Levels of free cysteine may fall during the massive inductions of cysteine-rich secreted proteins in development or immune defense, as well as during the synthesis of collagens in the periodic molting cycle (54, 80). The oxidation of so many cysteines to disulfides in the endoplasmic reticulum might locally lower O_2_ levels preventing EGL-9 hydroxylation of HIF-1. Thus, the intersection between HIF-1 and cysteine homeostasis we have uncovered may contribute to a regulatory axis in cell-cell signaling during development and in immune function.

## Methods

### General methods and strains

*C. elegans* were cultured using established protocols (43). Briefly, animals were cultured at 20°C on nematode growth media (NGM) seeded with wild-type *E. coli* (OP50). The wild-type strain of *C. elegans* was Bristol N2. Additional *E. coli* strains used in this work were BW25113 (Wild type, Moco+) and JW0764-2 (Δ*moaA753::kan,* Moco-) (81).

*C. elegans* mutant and transgenic strains used in this work are listed here. When previously published, sources of strains are referenced. Unless a reference is provided, all strains were generated in this study.

#### Non-transgenic *C. elegans*

ZG31, *hif-1(ia4) V* (82)
JT307, *egl-9(sa307) V* (44)
GR2254, *moc-1(ok366) X* (33)
GR2260, *cdo-1(mg622) X* (33)
GR2261, *cdo-1(mg622) moc-1(ok366) X* (33)
GR2269, *suox-1(gk738847) X* (33)
CB5602, *vhl-1(ok161) X* (39)
USD410, *cysl-1(ok762) X,* outcrossed 3x for this work
USD414, *rhy-1(ok1402) II; suox-1(gk738847) X*
USD421, *egl-9(sa307) V; suox-1(gk738847) X*
USD422, *vhl-1(ok161) suox-1(gk738847) X*
USD430, *egl-9(sa307) V; cdo-1(mg622) suox-1(gk738847) X*
USD431, *hif-1(ia4) egl-9(sa307) V; suox-1(gk738847) X*
USD432, *rhy-1(ok1402) II; egl-9(sa307) V; suox-1(gk738847) X*
USD433, *cth-2(mg599) II; egl-9(sa307) V; suox-1(gk738847) X*
USD434, *egl-9(sa307) X; cysl-1(ok762) suox-1(gk738847) X*
USD512, *rhy-1(ok1402) II,* outcrossed 4x for this work
USD706*, unc-119(ed3) III; cdo-1(mg622) moc-1(ok366) X*
USD920, *cdo-1(rae273) moc-1(ok366) X*
USD921, *egl-9(sa307) V; cdo-1(rae273) X*
USD922, *rhy-1(ok1402) II; cdo-1(rae273) X*
USD937, *egl-9(rae276) V; suox-1(gk738847) X*

#### MiniMos transgenic lines

USD531, *unc-119(ed3) III; raeTi1 [Pcdo-1::CDO-1::GFP unc-119(+)]*
USD719, *unc-119(ed3) III; cdo-1(mg622) moc-1(ok366); raeTi14 [Pcdo-1::CDO-1(C85Y)::GFP unc-119(+)]*
USD720, *unc-119(ed3) III; raeTi15 [Pcdo-1::GFP unc-119(+)]*
USD730, *rhy-1(ok1402) II; unc-119(ed3) III; raeTi15 [Pcdo-1::GFP unc-119(+)]*
USD733, *unc-119(ed3) III; egl-9(sa307) V; raeTi15 [Pcdo-1::GFP unc-119(+)]*
USD739, *unc-119(ed3) III; cdo-1(mg622) moc-1(ok366) X; raeTi1 [Pcdo-1::CDO-1::GFP unc-119(+)]*
USD766, *unc-119(ed3) III; cdo-1(mg622) moc-1(ok366) X; raeTi32 [Pcol-10::CDO-1::GFP unc-119(+)]*
USD767, *unc-119(ed3) III; cdo-1(mg622) moc-1(ok366) X; raeTi33 [Pcol-10::CDO-1::GFP unc-119(+)]*
USD776, *rhy-1(ok1402) II; unc-119(ed3) III; hif-1(ia4) V; raeTi15 [Pcdo-1::GFP unc-119(+)]*
USD777, *unc-119(ed3) III; egl-9(sa307) hif-1(ia4) V; raeTi15 [Pcdo-1::GFP unc-119(+)]*
USD780, *rhy-1(ok1402) II; unc-119(ed3) III; cysl-1(ok762) X; raeTi15 [Pcdo-1::GFP unc-119(+)]*
USD787, *unc-119(ed3) III; egl-9(sa307) V; cysl-1(ok762) X; raeTi15 [Pcdo-1::GFP unc-119(+)]*
USD808, *unc-119(ed3) III; cdo-1(mg622) moc-1(ok366) X; raeTi40 [Pcol-10::CDO-1[C85Y]::GFP unc-119(+)]*
USD810, *unc-119(ed3) III; cdo-1(mg622) moc-1(ok366) X; raeTi41 [Pcol-10::CDO-1[C85Y]::GFP unc-119(+)]*
USD940, *unc-119(ed3) III; vhl-1(ok161) X; raeTi15 [Pcdo-1::GFP unc-119(+)]*
USD1160, *unc-119(ed3) III; cysl-1(ok762) X; raeTi15 [Pcdo-1::GFP unc-119(+)]*
USD1161, *unc-119(ed3) III; hif-1(ia4) V; raeTi15 [Pcdo-1::GFP unc-119(+)]*

#### EMS-derived strains

USD659, *unc-119(ed3) III; egl-9(rae213) V; raeTi1 [Pcdo-1::CDO-1::GFP unc-119(+)]*
USD674, *unc-119(ed3) III; egl-9(rae227) V; raeTi1 [Pcdo-1::CDO-1::GFP unc-119(+)]*
USD655, *rhy-1(rae209) II; unc-119(ed3) III; raeTi1 [Pcdo-1::CDO-1::GFP unc-119(+)]*
USD656, *rhy-1(rae210) II; unc-119(ed3) III; raeTi1 [Pcdo-1::CDO-1::GFP unc-119(+)]*
USD657, *rhy-1(rae211) II; unc-119(ed3) III; raeTi1 [Pcdo-1::CDO-1::GFP unc-119(+)]*
USD658, *rhy-1(rae212) II; unc-119(ed3) III; raeTi1 [Pcdo-1::CDO-1::GFP unc-119(+)]*

#### CRISPR/Cas9-derived strains

USD914, *cdo-1(rae273) X,* CDO-1::GFP
USD926, *egl-9(rae276) V,* EGL-9[H487A]
USD928, *unc-119(ed3) III; egl-9(rae278) V; raeTi15 [Pcdo-1::GFP unc-119(+)]*

### MiniMos transgenesis

Cloning of original plasmid constructs was performed using isothermal/Gibson assembly (83). All MiniMos constructs were assembled in pNL43, which is derived from pCFJ909, a gift from Erik Jorgensen (Addgene plasmid #44480) (84). Details about plasmid construction are described below. MiniMos transgenic animals were generated using established protocols that rescue the *unc-119(ed3)* Unc phenotype (41).

To generate a construct that expressed CDO-1 under the control of its native promoter, we cloned the wild-type *cdo-1* genomic locus from 1,335 base pairs upstream of the *cdo-1* ATG start codon to (and including) codon 190 encoding the final CDO-1 amino acid prior to the TAA stop codon. This wild-type genomic sequence was fused in frame with a C-terminal GFP and *tbb-2* 3’UTR (376 bp downstream of the *tbb-2* stop codon) in the pNL43 plasmid backbone. This plasmid is called pKW24 (P*cdo-1::CDO-1::GFP*).

To generate pKW44, a construct encoding the active site mutant transgene *Pcdo-1::CDO-1(C85Y)::GFP,* we performed Q5 site-directed mutagenesis on pKW24, following manufacturer’s instructions (New England Biolabs). In pKW44, codon 85 was mutated from TGC (cysteine) to TAC (tyrosine).

To generate pKW45 (P*cdo-1::GFP*), the 1,335 base pair *cdo-1* promoter was amplified and fused directly to the GFP coding sequence. Both fragments were derived from pKW24, excluding the *cdo-1* coding sequence and introns.

pKW49 is a construct driving *cdo-1* expression from the hypodermal-specific *col-10* promoter (*Pcol-10::CDO-1::GFP*) (53). The *col-10* promoter (1,126 base pairs upstream of the *col-10* start codon) was amplified and fused upstream of the *cdo-1* ATG start codon in pKW24, replacing the native *cdo-1* promoter. pKW53 [P*col-10::CDO-1(C85Y)::GFP*] was engineered using the same Q5-site-directed mutagenesis strategy as was described for pKW44. However, this mutagenesis used pKW49 as the template plasmid.

### Chemical mutagenesis and whole genome sequencing

To define *C. elegans* gene activities that were necessary for the control of *cdo-1* levels, we carried out a chemical mutagenesis screen for animals that accumulate CDO-1 protein. To visualize CDO-1 levels, we engineered USD531, a transgenic *C. elegans* strain carrying the *raeTi1* [pKW24, *Pcdo-1::CDO-1::GFP*] transgene. USD531 transgenic *C. elegans* were mutagenized with ethyl methanesulfonate (EMS) using established protocols (43). F2 generation animals were manually screened, and mutant isolates were collected that displayed high accumulation of *Pcdo-1::CDO-1::GFP.* We demanded that mutant strains of interest were viable and fertile.

We followed established protocols to identify EMS-induced mutations in our strains of interest (85). Briefly, whole genomic DNA was prepared from *C. elegans* using the Gentra Puregene Tissue Kit (Qiagen) and genomic DNA libraries were prepared using the NEBNext genomic DNA library construction kit (New England Biolabs). DNA libraries were sequenced on an Illumina Hi-Seq and deep sequencing reads were analyzed using standard methods on Galaxy, a web-based platform for computational analyses (86). Briefly, sequencing reads were trimmed and aligned to the WBcel235 *C. elegans* reference genome (87, 88). Variations from the reference genome and the putative impact of those variations were annotated and extracted for analysis (89–91). Here we report the analysis of 6 new mutant strains (USD655, USD656, USD657, USD658, USD659, and USD674). Among these mutant strains, we found 2 unique mutations in *egl-9* (USD659 and USD674) and 4 unique mutations in *rhy-1* (USD655, USD656, USD657, USD658). The allele names and molecular identity of these new *egl-9* and *rhy-1* mutations are specified in Fig. 1C. These genes were prioritized based on the isolation of multiple independent alleles and their established functions in a common pathway, the hypoxia and H_2_S-sensing pathway (36–38).

### Genome engineering by CRISPR/Cas9

We followed standard protocols to perform CRISPR/Cas9 genome engineering of *cdo-1* and *egl-9* genomic loci in *C. elegans* (92–95). Essential details of the CRISPR/Cas9-generated reagents in this work are described below.

We used homology-directed repair to generate *cdo-1(rae273)* [CDO-1::GFP]. The guide RNA was 5’-gactacagaggatctaagaa-3’ (crRNA, IDT). The GFP donor double-stranded DNA (dsDNA) was amplified from pCFJ2249 using primers that contained roughly 40bp of homology to *cdo-1* flanking the desired insertion site (96). The primers used to generate the donor dsDNA were: 5’-gtacggcaagaaagttgactacagaggatctaagaataatagtactagcggtggcagtgg-3’ and 5’-agaatcaacacgttattacattgagggatatgttgtttacttgtagagctcgtccattcc-3’. Glycine 189 of CDO-1 was removed by design to eliminate the PAM site and prevent cleavage of the donor dsDNA.

The *egl-9(rae276)* and *egl-9(rae278)* [EGL-9(H487A)] alleles were also generated by homology-directed repair using the same combination of guide RNA and single-stranded oligodeoxynucleotide (ssODN) donor. The guide RNA was 5’-tgtgaagcatgtagataatc-3’ (crRNA, IDT) The ssODN donor was 5’-gcttgccatctatcctggaaatggaactcgttatgtgaaggctgtagacaatccagtaaaagatggaagatgtataaccactatttattactg-3’ (Ultramer, IDT). Successful editing resulted in altering the coding sequence of EGL-9 to encode for an alanine rather than the catalytically essential histidine at position 487. We also used synonymous mutations to introduce an AccI restriction site that is helpful for genotyping the *rae276* and *rae278* mutant alleles.

### *C. elegans* growth assays

To assay developmental rates, *C. elegans* were synchronized at the first stage of larval development. To synchronize animals, embryos were harvested from gravid adult animals via treatment with a bleach and sodium hydroxide solution. Embryos were then incubated overnight in M9 solution causing them to hatch and arrest development at the L1 stage (97). Synchronized L1 animals were cultured on NGM seeded with BW25113 (Wild type, Moco+) or JW0764-2 (Δ*moaA753::kan,* Moco-) *E. coli.* Animals were cultured for 48 or 72 hours (specified in the appropriate figure legends), and live animals were imaged as described below. Animal length was measured from tip of head to the end of the tail.

To determine qualitative ‘health’ of various *C. elegans* strains, we assayed the ability of these strains to consume all *E. coli* food provided on an NGM petri dish. For this experiment, dietary *E. coli* was produced via overnight culture in liquid LB in a 37°C shaking incubator. 200μl of this *E. coli* was seeded onto NGM petri dishes and allowed to dry, producing nearly identical lawns and growth environments. Then, 5 L4 animals of a strain of interest were introduced onto these NGM petri dishes seeded with OP50. For each experiment, petri dishes were monitored daily and scored when all *E. coli* was consumed by the population of animals. This assay is beneficial because it integrates many life-history measures (i.e. developmental rate, brood size, embryonic viability, etc.) into a single simple assay that can be scaled and applied to many *C. elegans* strains in parallel.

### Cysteine exposure

To determine the impact of supplemental cysteine on expression of *cdo-1* and animal viability, we exposed various *C. elegans* strains to 0, 50, 100, 250, 500, or 1,000μM supplemental cysteine. *C. elegans* strains were synchronously grown on NGM media supplemented with *E. coli* OP50 at 20°C until reaching the L4 stage of development. Live L4 animals were then collected from the petri dishes and cultured in liquid M9 media containing 4X concentrated *E. coli* OP50 with or without supplemental cysteine. These liquid cultures were gently rocked at 20°C overnight. Post exposure, GFP imaging was performed as described in the Microscopy section of the materials and methods. Alternatively, animals exposed overnight to 0, 100, or 1,000μM supplemental cysteine were scored for viability after being seeded onto NGM petri dishes. Animals were determined to be alive if they responded to mechanical stimulus.

### Microscopy

Low magnification bright field and fluorescence images (imaging GFP simultaneously in multiple animals) were collected using a Zeiss AxioZoom V16 microscope equipped with a Hamamatsu Orca flash 4.0 digital camera using Zen software (Zeiss). For experiments with supplemental cysteine, low magnification bright field and fluorescence images were collected using a Nikon SMZ25 microscope equipped with a Hamamatsu Orca flash 4.0 digital camera using NIS-Elements software (Nikon). High magnification differential interference contrast (DIC) and GFP fluorescence images (imaging CDO-1::GFP encoded by *cdo-1(rae273)*) were collected using Zeiss AxioImager Z1 microscope equipped with a Zeiss AxioCam HRc digital camera using Zen software (Zeiss). All images were processed and analyzed using ImageJ software (NIH). All imaging was performed on live animals paralyzed using 20mM sodium azide. For all fluorescence images shown within the same figure panel, images were collected using the same exposure time and processed identically. To quantify GFP expression, the average pixel intensity was determined within a set transverse section immediately anterior to the developing vulva. Background pixel intensity was determined in a set region of interest distinct from the *C. elegans* samples and was subtracted from the sample measurements.

## Acknowledgements

Some *C. elegans* strains were provided by the CGC, which is funded by the NIH Office of Research Infrastructure Programs (P40 OD010440). Research reported in this publication was supported by the National Institute of General Medical Sciences of the National Institutes of Health under award numbers R01 GM044619 (to G.R.) and R35 GM146871 (to K.W.).

## Supplemental Figures

**Figure S1:**
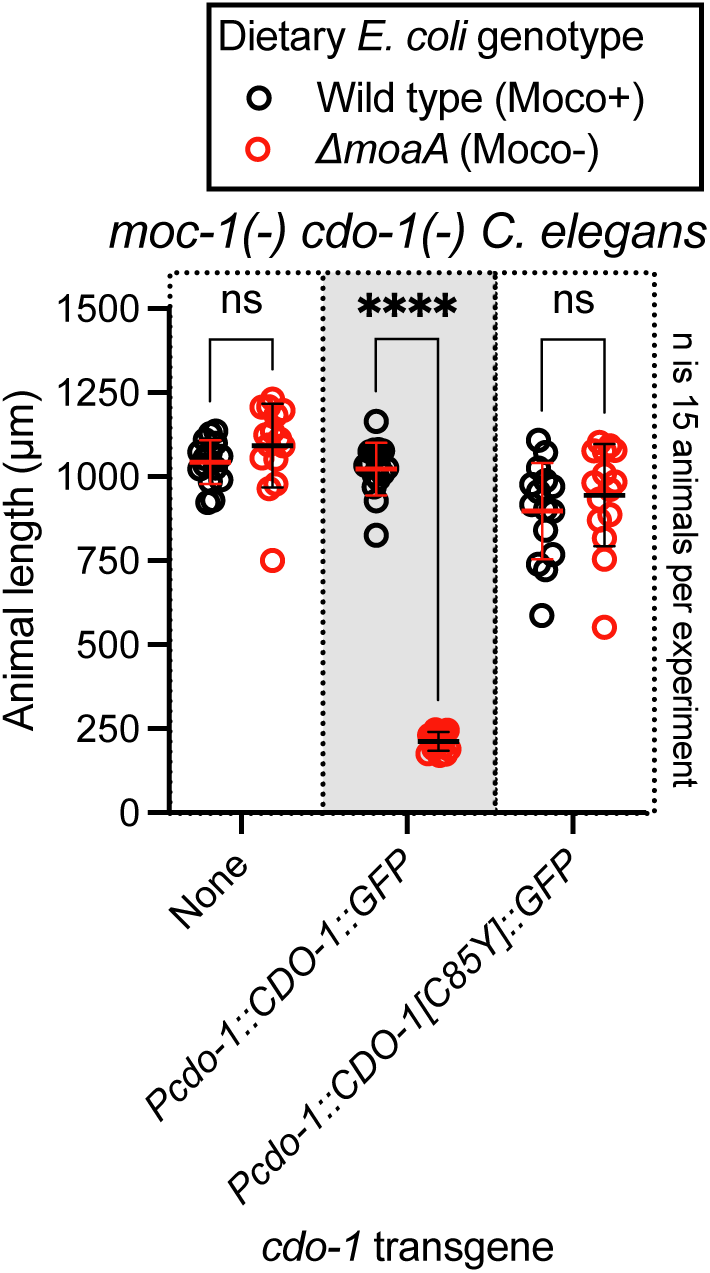
The *Pcdo-1::CDO-1::GFP* transgene encodes a functional cysteine dioxygenase enzyme. *moc-1(ok366) cdo-1(mg622)* double mutant animals expressing *Pcdo-1::CDO-1::GFP* or *Pcdo-1::CDO-1[C85Y]::GFP* transgenes were cultured from synchronized L1 larvae for 72 hours on wild-type (black, Moco+) or Δ*moaA* mutant (red, Moco-) *E. coli.* Animal lengths were determined for each condition. Individual datapoints are shown (circles) as are the mean and standard deviation. Sample size (*n*) is displayed for each experiment. ****, p<0.0001, multiple unpaired t test with Welch’s correction. ns indicates no significant difference was identified.

**Figure S2:**
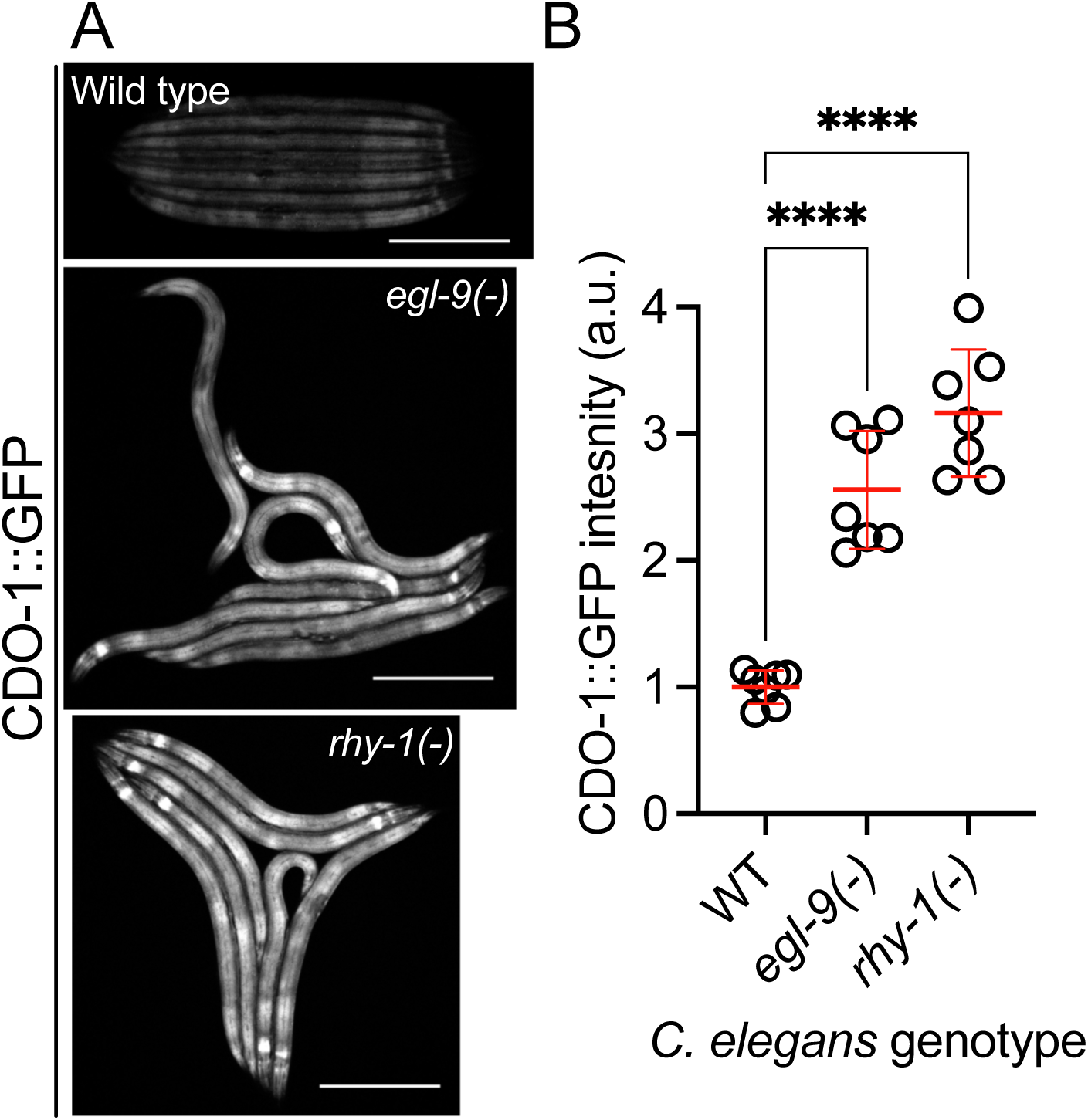
A functional CDO-1::GFP fusion protein is induced by loss of *egl-9* or *rhy-1*. A) Expression of CDO-1::GFP from the *cdo-1(rae273)* allele is displayed for wild-type, *egl-9(sa307),* and *rhy-1(ok1402)* animals at the L4 stage of development. Scale bar is 250μm. For GFP imaging, exposure time was 100ms. B) Quantification of CDO-1::GFP expression displayed in Fig. S2A. Individual datapoints are shown (circles) as are the mean and standard deviation (red lines). *n* is 7 individuals per genotype. Data are normalized so that wild-type expression of CDO-1::GFP is 1 arbitrary unit (a.u.). ****, p<0.0001, ordinary one-way ANOVA with Dunnett’s post hoc analysis.

**Figure S3:**
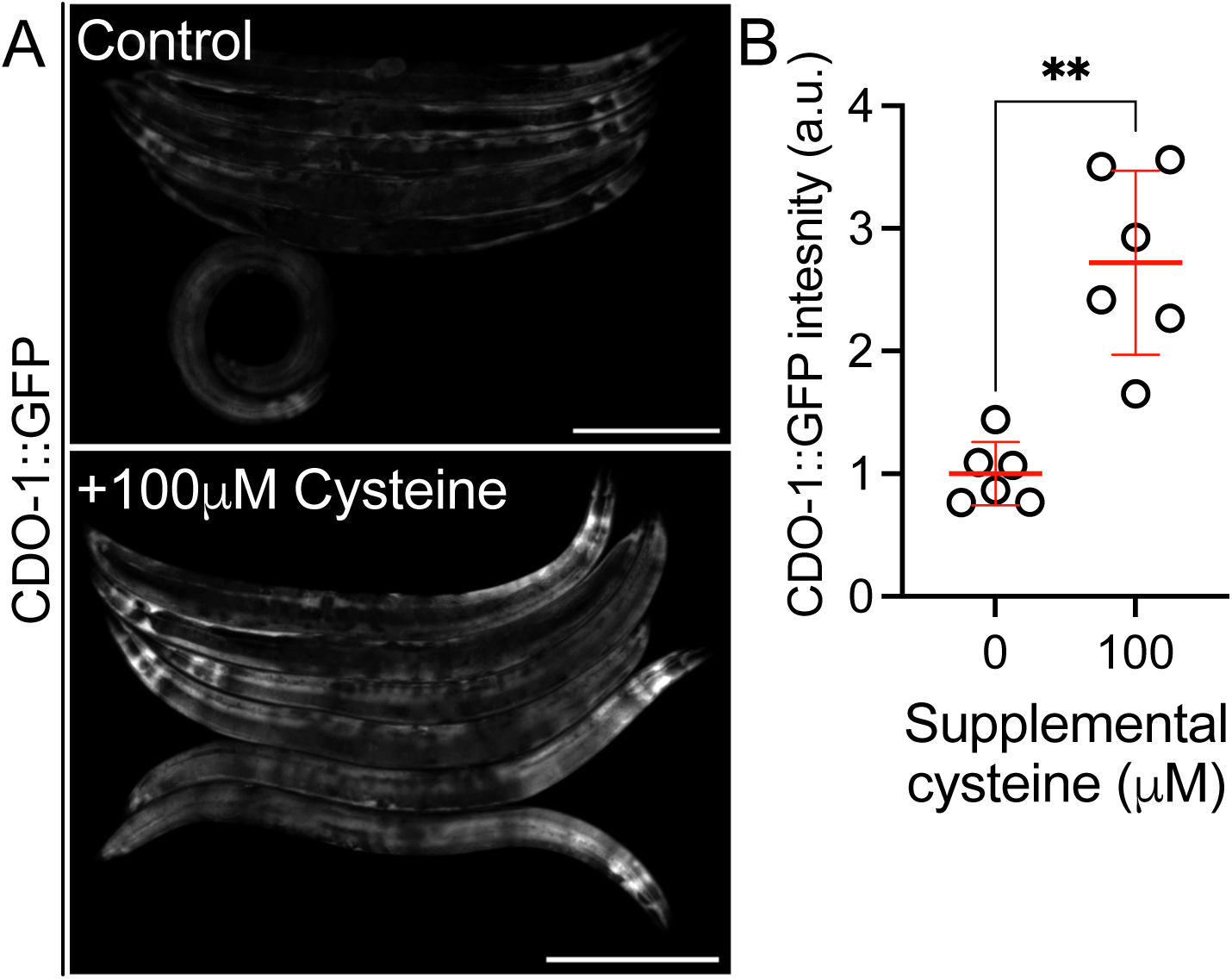
CDO-1::GFP encoded by *cdo-1(rae273)* is induced by supplemental cysteine. A) Expression of CDO-1::GFP from the *cdo-1(rae273)* allele is displayed for wild-type young adult animals exposed to 0 or 100μM supplemental cysteine. Scale bar is 250μm. For GFP imaging, exposure time was 100ms. Supplemental cysteine did not impact the mortality of the animals being imaged. B) Quantification of CDO-1::GFP expression displayed in Fig. S3A. Individual datapoints are shown (circles) as are the mean and standard deviation (red lines). *n* is 6 individuals per genotype. Data are normalized so that wild-type expression of CDO-1::GFP exposed to 0μM supplemental cysteine is 1 arbitrary unit (a.u.). **, p<0.001, unpaired t test with Welch’s correction.

## Notes

### Competing Interest Statement

The authors have declared no competing interest.

### Summary of Updates

This version of the manuscript has been updated to add new data to Figure 3, modify existing figures, and include additional comments to the Results and Discussion.

## References

1. G. Wu, Y. Z. Fang, S. Yang, J. R. Lupton, N. D. Turner, Glutathione metabolism and its implications for health. J Nutr 134, 489–492 (2004).

2. L. Zheng, R. H. White, V. L. Cash, R. F. Jack, D. R. Dean, Cysteine desulfurase activity indicates a role for NIFS in metallocluster biosynthesis. Proc Natl Acad Sci U S A 90, 2754–2758 (1993).

3. R. Noiva, Enzymatic catalysis of disulfide formation. Protein Expr Purif 5, 1–13 (1994).

4. S. Raina, D. Missiakas, Making and breaking disulfide bonds. Annu Rev Microbiol 51, 179–202 (1997).

5. A. Rietsch, J. Beckwith, The genetics of disulfide bond metabolism. Annu Rev Genet 32, 163–184 (1998).

6. N. M. Giles et al., Metal and redox modulation of cysteine protein function. Chem Biol 10, 677–693 (2003).

7. J. A. Tainer, V. A. Roberts, E. D. Getzoff, Metal-binding sites in proteins. Curr Opin Biotechnol 2, 582–591 (1991).

8. J. Miller, A. D. McLachlan, A. Klug, Repetitive zinc-binding domains in the protein transcription factor IIIA from Xenopus oocytes. Embo j 4, 1609–1614 (1985).

9. S. Singh, R. Banerjee, PLP-dependent H(2)S biogenesis. Biochim Biophys Acta 1814, 1518–1527 (2011).

10. C. E. Hughes et al., Cysteine Toxicity Drives Age-Related Mitochondrial Decline by Altering Iron Homeostasis. Cell 180, 296–310.e218 (2020).

11. J. W. Olney, C. Zorumski, M. T. Price, J. Labruyere, L-cysteine, a bicarbonate-sensitive endogenous excitotoxin. Science 248, 596–599 (1990).

12. C. L. Evans, The toxicity of hydrogen sulphide and other sulphides. Q J Exp Physiol Cogn Med Sci 52, 231–248 (1967).

13. D. H. Truong, M. A. Eghbal, W. Hindmarsh, S. H. Roth, P. J. O’Brien, Molecular mechanisms of hydrogen sulfide toxicity. Drug Metab Rev 38, 733–744 (2006).

14. D. L. Bella, Y. H. Kwon, M. H. Stipanuk, Variations in dietary protein but not in dietary fat plus cellulose or carbohydrate levels affect cysteine metabolism in rat isolated hepatocytes. J Nutr 126, 2179–2187 (1996).

15. M. H. Stipanuk, Sulfur amino acid metabolism: pathways for production and removal of homocysteine and cysteine. Annu Rev Nutr 24, 539–577 (2004).

16. M. H. Stipanuk, I. Ueki, Dealing with methionine/homocysteine sulfur: cysteine metabolism to taurine and inorganic sulfur. J Inherit Metab Dis 34, 17–32 (2011).

17. J. E. Dominy, Jr., L. L. Hirschberger, R. M. Coloso, M. H. Stipanuk, In vivo regulation of cysteine dioxygenase via the ubiquitin-26S proteasome system. Adv Exp Med Biol 583, 37–47 (2006).

18. J. E. Dominy, Jr., L. L. Hirschberger, R. M. Coloso, M. H. Stipanuk, Regulation of cysteine dioxygenase degradation is mediated by intracellular cysteine levels and the ubiquitin-26 S proteasome system in the living rat. Biochem J 394, 267–273 (2006).

19. M. H. Stipanuk, L. L. Hirschberger, M. P. Londono, C. L. Cresenzi, A. F. Yu, The ubiquitin-proteasome system is responsible for cysteine-responsive regulation of cysteine dioxygenase concentration in liver. Am J Physiol Endocrinol Metab 286, E439–448 (2004).

20. J. I. Lee, M. Londono, L. L. Hirschberger, M. H. Stipanuk, Regulation of cysteine dioxygenase and gamma-glutamylcysteine synthetase is associated with hepatic cysteine level. J Nutr Biochem 15, 112–122 (2004).

21. Y. H. Kwon, M. H. Stipanuk, Cysteine regulates expression of cysteine dioxygenase and gamma-glutamylcysteine synthetase in cultured rat hepatocytes. Am J Physiol Endocrinol Metab 280, E804–815 (2001).

22. K. Kojima et al., Cysteine dioxygenase type 1 (CDO1) gene promoter methylation during the adenoma-carcinoma sequence in colorectal cancer. PLoS One 13, e0194785 (2018).

23. K. Yamashita, K. Hosoda, N. Nishizawa, H. Katoh, M. Watanabe, Epigenetic biomarkers of promoter DNA methylation in the new era of cancer treatment. Cancer Sci 109, 3695–3706 (2018).

24. M. Brait et al., Cysteine dioxygenase 1 is a tumor suppressor gene silenced by promoter methylation in multiple human cancers. PLoS One 7, e44951 (2012).

25. Y. P. Kang et al., Cysteine dioxygenase 1 is a metabolic liability for non-small cell lung cancer. Elife 8 (2019).

26. J. Jeschke et al., Frequent inactivation of cysteine dioxygenase type 1 contributes to survival of breast cancer cells and resistance to anthracyclines. Clin Cancer Res 19, 3201–3211 (2013).

27. S. Hao et al., Cysteine Dioxygenase 1 Mediates Erastin-Induced Ferroptosis in Human Gastric Cancer Cells. Neoplasia 19, 1022–1032 (2017).

28. G. Ma et al., Cysteine dioxygenase 1 attenuates the proliferation via inducing oxidative stress and integrated stress response in gastric cancer cells. Cell Death Discov 8, 493 (2022).

29. M. Duran et al., Combined deficiency of xanthine oxidase and sulphite oxidase: a defect of molybdenum metabolism or transport? J Inherit Metab Dis 1, 175–178 (1978).

30. S. H. Mudd, F. Irreverre, L. Laster, Sulfite oxidase deficiency in man: demonstration of the enzymatic defect. Science 156, 1599–1602 (1967).

31. G. Schwarz, R. R. Mendel, M. W. Ribbe, Molybdenum cofactors, enzymes and pathways. Nature 460, 839–847 (2009).

32. Y. Zhang, V. N. Gladyshev, Molybdoproteomes and evolution of molybdenum utilization. J Mol Biol 379, 881–899 (2008).

33. K. Warnhoff, G. Ruvkun, Molybdenum cofactor transfer from bacteria to nematode mediates sulfite detoxification. Nat Chem Biol 15, 480–488 (2019).

34. K. Warnhoff, T. W. Hercher, R. R. Mendel, G. Ruvkun, Protein-bound molybdenum cofactor is bioavailable and rescues molybdenum cofactor-deficient C. elegans. Genes Dev 35, 212–217 (2021).

35. J. Snoozy, P. C. Breen, G. Ruvkun, K. Warnhoff, moc-6/MOCS2A is necessary for molybdenum cofactor synthesis in C. elegans. MicroPubl Biol 2022 (2022).

36. C. Shen, Z. Shao, J. A. Powell-Coffman, The Caenorhabditis elegans rhy-1 gene inhibits HIF-1 hypoxia-inducible factor activity in a negative feedback loop that does not include vhl-1. Genetics 174, 1205–1214 (2006).

37. M. W. Budde, M. B. Roth, The response of Caenorhabditis elegans to hydrogen sulfide and hydrogen cyanide. Genetics 189, 521–532 (2011).

38. D. K. Ma, R. Vozdek, N. Bhatla, H. R. Horvitz, CYSL-1 interacts with the O2-sensing hydroxylase EGL-9 to promote H2S-modulated hypoxia-induced behavioral plasticity in C. elegans. Neuron 73, 925–940 (2012).

39. A. C. Epstein et al., C. elegans EGL-9 and mammalian homologs define a family of dioxygenases that regulate HIF by prolyl hydroxylation. Cell 107, 43–54 (2001).

40. M. Ivan et al., HIFalpha targeted for VHL-mediated destruction by proline hydroxylation: implications for O2 sensing. Science 292, 464–468 (2001).

41. C. Frøkjær-Jensen et al., Random and targeted transgene insertion in Caenorhabditis elegans using a modified Mos1 transposon. Nat Methods 11, 529–534 (2014).

42. J. G. McCoy et al., Structure and mechanism of mouse cysteine dioxygenase. Proc Natl Acad Sci U S A 103, 3084–3089 (2006).

43. S. Brenner, The genetics of Caenorhabditis elegans. Genetics 77, 71–94 (1974).

44. C. Darby, C. L. Cosma, J. H. Thomas, C. Manoil, Lethal paralysis of Caenorhabditis elegans by Pseudomonas aeruginosa. Proc Natl Acad Sci U S A 96, 15202–15207 (1999).

45. B. Wang et al., Co-opted genes of algal origin protect C. elegans against cyanogenic toxins. Curr Biol 32, 4941–4948.e4943 (2022).

46. S. Salceda, J. Caro, Hypoxia-inducible factor 1alpha (HIF-1alpha) protein is rapidly degraded by the ubiquitin-proteasome system under normoxic conditions. Its stabilization by hypoxia depends on redox-induced changes. J Biol Chem 272, 22642–22647 (1997).

47. L. E. Huang, J. Gu, M. Schau, H. F. Bunn, Regulation of hypoxia-inducible factor 1alpha is mediated by an O2-dependent degradation domain via the ubiquitin-proteasome pathway. Proc Natl Acad Sci U S A 95, 7987–7992 (1998).

48. C. H. Sutter, E. Laughner, G. L. Semenza, Hypoxia-inducible factor 1alpha protein expression is controlled by oxygen-regulated ubiquitination that is disrupted by deletions and missense mutations. Proc Natl Acad Sci U S A 97, 4748–4753 (2000).

49. C. L. Pender, H. R. Horvitz, Hypoxia-inducible factor cell non-autonomously regulates C. elegans stress responses and behavior via a nuclear receptor. Elife 7 (2018).

50. M. M. Kudron et al., The ModERN Resource: Genome-Wide Binding Profiles for Hundreds of Drosophila and Caenorhabditis elegans Transcription Factors. Genetics 208, 937–949 (2018).

51. M. Vora et al., The hypoxia response pathway promotes PEP carboxykinase and gluconeogenesis in C. elegans. Nat Commun 13, 6168 (2022).

52. W. G. Kaelin, Jr., P. J. Ratcliffe, Oxygen sensing by metazoans: the central role of the HIF hydroxylase pathway. Mol Cell 30, 393–402 (2008).

53. Y. Hong, R. C. Lee, V. Ambros, Structure and function analysis of LIN-14, a temporal regulator of postembryonic developmental events in Caenorhabditis elegans. Mol Cell Biol 20, 2285–2295 (2000).

54. M. W. Meeuse et al., Developmental function and state transitions of a gene expression oscillator in Caenorhabditis elegans. Mol Syst Biol 16, e9498 (2020).

55. K. D. Oliphant, R. R. Fettig, J. Snoozy, R. R. Mendel, K. Warnhoff, Obtaining the necessary molybdenum cofactor for sulfite oxidase activity in the nematode Caenorhabditis elegans surprisingly involves a dietary source. J Biol Chem 299, 102736 (2023).

56. J. W. Horsman, F. I. Heinis, D. L. Miller, A Novel Mechanism To Prevent H(2)S Toxicity in Caenorhabditis elegans. Genetics 213, 481–490 (2019).

57. Y. Pan et al., Multiple factors affecting cellular redox status and energy metabolism modulate hypoxia-inducible factor prolyl hydroxylase activity in vivo and in vitro. Mol Cell Biol 27, 912–925 (2007).

58. Z. Shao, Y. Zhang, J. A. Powell-Coffman, Two distinct roles for EGL-9 in the regulation of HIF-1-mediated gene expression in Caenorhabditis elegans. Genetics 183, 821–829 (2009).

59. G. L. Semenza, G. L. Wang, A nuclear factor induced by hypoxia via de novo protein synthesis binds to the human erythropoietin gene enhancer at a site required for transcriptional activation. Mol Cell Biol 12, 5447–5454 (1992).

60. N. Bashan, E. Burdett, H. S. Hundal, A. Klip, Regulation of glucose transport and GLUT1 glucose transporter expression by O2 in muscle cells in culture. Am J Physiol 262, C682–690 (1992).

61. J. D. Loike et al., Hypoxia induces glucose transporter expression in endothelial cells. Am J Physiol 263, C326–333 (1992).

62. J. D. Firth, B. L. Ebert, C. W. Pugh, P. J. Ratcliffe, Oxygen-regulated control elements in the phosphoglycerate kinase 1 and lactate dehydrogenase A genes: similarities with the erythropoietin 3’ enhancer. Proc Natl Acad Sci U S A 91, 6496–6500 (1994).

63. J. M. Gray et al., Oxygen sensation and social feeding mediated by a C. elegans guanylate cyclase homologue. Nature 430, 317–322 (2004).

64. M. Hirsilä, P. Koivunen, V. Günzler, K. I. Kivirikko, J. Myllyharju, Characterization of the human prolyl 4-hydroxylases that modify the hypoxia-inducible factor. J Biol Chem 278, 30772–30780 (2003).

65. J. H. Dao et al., Kinetic characterization and identification of a novel inhibitor of hypoxia-inducible factor prolyl hydroxylase 2 using a time-resolved fluorescence resonance energy transfer-based assay technology. Anal Biochem 384, 213–223 (2009).

66. D. Ehrismann et al., Studies on the activity of the hypoxia-inducible-factor hydroxylases using an oxygen consumption assay. Biochem J 401, 227–234 (2007).

67. L. Li et al., Searching for molecular hypoxia sensors among oxygen-dependent enzymes. Elife 12 (2023).

68. J. A. Losman, P. Koivunen, W. G. Kaelin, Jr., 2-Oxoglutarate-dependent dioxygenases in cancer. Nat Rev Cancer 20, 710–726 (2020).

69. M. W. Budde, M. B. Roth, Hydrogen sulfide increases hypoxia-inducible factor-1 activity independently of von Hippel-Lindau tumor suppressor-1 in C. elegans. Mol Biol Cell 21, 212–217 (2010).

70. D. L. Miller, M. W. Budde, M. B. Roth, HIF-1 and SKN-1 coordinate the transcriptional response to hydrogen sulfide in Caenorhabditis elegans. PLoS One 6, e25476 (2011).

71. C. Shen, D. Nettleton, M. Jiang, S. K. Kim, J. A. Powell-Coffman, Roles of the HIF-1 hypoxia-inducible factor during hypoxia response in Caenorhabditis elegans. J Biol Chem 280, 20580–20588 (2005).

72. H. Jurkowska et al., Primary hepatocytes from mice lacking cysteine dioxygenase show increased cysteine concentrations and higher rates of metabolism of cysteine to hydrogen sulfide and thiosulfate. Amino Acids 46, 1353–1365 (2014).

73. Y. J. Peng et al., H2S mediates O2 sensing in the carotid body. Proc Natl Acad Sci U S A 107, 10719–10724 (2010).

74. K. J. Briggs et al., Paracrine Induction of HIF by Glutamate in Breast Cancer: EglN1 Senses Cysteine. Cell 166, 126–139 (2016).

75. S. Bannai, E. Kitamura, Transport interaction of L-cystine and L-glutamate in human diploid fibroblasts in culture. J Biol Chem 255, 2372–2376 (1980).

76. H. Sato, M. Tamba, T. Ishii, S. Bannai, Cloning and expression of a plasma membrane cystine/glutamate exchange transporter composed of two distinct proteins. J Biol Chem 274, 11455–11458 (1999).

77. N. O. Burton et al., Intergenerational adaptations to stress are evolutionarily conserved, stress-specific, and have deleterious trade-offs. Elife 10 (2021).

78. G. M. Gibbs, K. Roelants, M. K. O’Bryan, The CAP superfamily: cysteine-rich secretory proteins, antigen 5, and pathogenesis-related 1 proteins--roles in reproduction, cancer, and immune defense. Endocr Rev 29, 865–897 (2008).

79. A. R. Frand, J. W. Cuozzo, C. A. Kaiser, Pathways for protein disulphide bond formation. Trends Cell Biol 10, 203–210 (2000).

80. N. Mizuki, M. Kasahara, Mouse submandibular glands express an androgen-regulated transcript encoding an acidic epididymal glycoprotein-like molecule. Mol Cell Endocrinol 89, 25–32 (1992).

81. T. Baba et al., Construction of Escherichia coli K-12 in-frame, single-gene knockout mutants: the Keio collection. Mol Syst Biol 2, 2006.0008 (2006).

82. H. Jiang, R. Guo, J. A. Powell-Coffman, The Caenorhabditis elegans hif-1 gene encodes a bHLH-PAS protein that is required for adaptation to hypoxia. Proc Natl Acad Sci U S A 98, 7916–7921 (2001).

83. D. G. Gibson et al., Enzymatic assembly of DNA molecules up to several hundred kilobases. Nat Methods 6, 343–345 (2009).

84. N. J. Lehrbach, G. Ruvkun, Proteasome dysfunction triggers activation of SKN-1A/Nrf1 by the aspartic protease DDI-1. Elife 5 (2016).

85. N. J. Lehrbach, F. Ji, R. Sadreyev, Next-Generation Sequencing for Identification of EMS-Induced Mutations in Caenorhabditis elegans. Curr Protoc Mol Biol 117, 7.29.21–27.29.12 (2017).

86. G. Community, The Galaxy platform for accessible, reproducible and collaborative biomedical analyses: 2022 update. Nucleic Acids Res 50, W345–351 (2022).

87. A. M. Bolger, M. Lohse, B. Usadel, Trimmomatic: a flexible trimmer for Illumina sequence data. Bioinformatics 30, 2114–2120 (2014).

88. H. Li, R. Durbin, Fast and accurate long-read alignment with Burrows-Wheeler transform. Bioinformatics 26, 589–595 (2010).

89. A. Wilm et al., LoFreq: a sequence-quality aware, ultra-sensitive variant caller for uncovering cell-population heterogeneity from high-throughput sequencing datasets. Nucleic Acids Res 40, 11189–11201 (2012).

90. P. Cingolani et al., Using Drosophila melanogaster as a Model for Genotoxic Chemical Mutational Studies with a New Program, SnpSift. Front Genet 3, 35 (2012).

91. P. Cingolani et al., A program for annotating and predicting the effects of single nucleotide polymorphisms, SnpEff: SNPs in the genome of Drosophila melanogaster strain w1118; iso-2; iso-3. Fly (Austin) 6, 80–92 (2012).

92. K. S. Ghanta, C. C. Mello, Melting dsDNA Donor Molecules Greatly Improves Precision Genome Editing in Caenorhabditis elegans. Genetics 216, 643–650 (2020).

93. J. A. Arribere et al., Efficient marker-free recovery of custom genetic modifications with CRISPR/Cas9 in Caenorhabditis elegans. Genetics 198, 837–846 (2014).

94. A. Paix, A. Folkmann, G. Seydoux, Precision genome editing using CRISPR-Cas9 and linear repair templates in C. elegans. Methods 121-122, 86–93 (2017).

95. G. A. Dokshin, K. S. Ghanta, K. M. Piscopo, C. C. Mello, Robust Genome Editing with Short Single-Stranded and Long, Partially Single-Stranded DNA Donors in Caenorhabditis elegans. Genetics 210, 781–787 (2018).

96. M. D. Aljohani, S. El Mouridi, M. Priyadarshini, A. M. Vargas-Velazquez, C. Frøkjær-Jensen, Engineering rules that minimize germline silencing of transgenes in simple extrachromosomal arrays in C. elegans. Nat Commun 11, 6300 (2020).

97. T. Stiernagle, Maintenance of C. elegans. WormBook 10.1895/wormbook.1.101.1, 1–11 (2006).

